# Identification of a novel subfamily of bacterial AAT-fold basic amino acid decarboxylases and functional characterization of its first representative: *Pseudomonas aeruginosa* LdcA

**DOI:** 10.1101/308080

**Authors:** Diego Carriel-Lopez, Pierre Simon Garcia, Florence Castelli, Patricia Lamourette, François Fenaille, Céline Brochier-Armanet, Sylvie Elsen, Irina Gutsche

## Abstract

Polyamines are small amino-acid derived polycations capable of binding negatively charged macromolecules. Bacterial polyamines are structurally and functionally diverse, and are mainly produced biosynthetically by PLP-dependent amino acid decarboxylases referred to as LAOdcs (Lysine-Arginine-Ornithine decarboxylases). In a phylogenetically limited group of bacteria, LAOdcs are also induced in response to acid stress. Here, we performed an exhaustive phylogenetic analysis of the AAT-fold LAOdcs which showcased the ancestral nature of their short forms in *Cyanobacteria* and *Firmicutes,* and emergence of distinct subfamilies of long LAOdcs in *Proteobacteria.* We identified a novel subfamily of lysine decarboxylases, LdcA, ancestral in *Betaproteobacteria* and *Pseudomortadaceae {Gammaproteobacteria).* We analyzed the expression of LdcA from *Pseudomonas aeruginosa,* and uncovered its role, intimately linked to cadaverine production, in promoting growth and reducing persistence of this multidrug resistant human pathogen during carbenicillin treatment. Finally, we documented a certain redundancy in the function of the three main polyamines - cadaverine, putrescine and spermidine - in *P. aeruginosa* by demonstrating the link between their intracellular level, as well as the capacity of putrescine and spermidine to complement the growth phenotype of the *ldcA* mutant.

## Introduction

Polyamines are small amino acid-derived molecules with two or more amino groups separated by alkyl chains (Tabor & Tabor, 1964, Lightfoot & Hall, 2014). They perform essential functions in all living organisms by participating in DNA replication, gene expression and protein synthesis, and are generally described as growth factors (Lightfoot & Hall, 2014, Michael, 2016b). At physiological pH, polyamines behave as polycations and can interact with negatively charged macromolecules such as nucleic acids, membrane phospholipids and proteins (Tabor & Tabor, 1964, Tabor & Tabor, 1985). They were proposed to bind and structurally modify RNA thereby acting at the level of translation (Igarashi & Kashiwagi, 2006, Igarashi & Kashiwagi, 2015). Strengthening this hypothesis, “polyamine modulons” were identified in *Escherichia coli* and more recently in eukaryotes (Igarashi & Kashiwagi, 2006, Igarashi & Kashiwagi, 2015). As a possible consequence, bacterial polyamines were shown to participate in expression of proteins essential for growth fitness and viability but also in processes such as biofilm formation, antibiotic resistance and virulence (Kwon & Lu, 2006, Shah & Swiatlo, 2008, Karatan & Michael, 2013, Di Martino *et al.,* 2013, Michael, 2016a).

The triamine spermidine (Spd), essential in eukaryotes and archaea (Lightfoot & Hall, 2014, Michael, 2016b), and its diamine precursor putrescine (Put) are extensively studied because these two polyamines are the only ones produced in all eukaryotes (Michael, 2016b). In bacteria, the polyamine repertoire and hence their roles are the most diverse. In particular, a third widespread bacterial polyamine, cadaverine (Cad), appears increasingly important for the physiology of *Proteobacteria.* Cad is recognized as an important player in enterobacterial acid stress response wherein it decreases porin permeability to protons and alkalinizes the medium (Dela Vega & Delcour, 1996, Zhao & Houry, 2010). During oxidative stress response, Cad is also capable of scavenging reactive oxygen species (ROS) in *Vibrio vulnificus* (Kang *et al.,* 2007). In some Negativicutes, *Selenomonas ruminantium* and *Veillonella sp.,* Cad was shown to be incorporated in the peptidoglycan and essential for its stability (Kamio *et al.,* 1986, Kamio & Nakamura, 1987). Finally, this polyamine is involved in iron uptake and is required for the synthesis of hydroxamate-type siderophores in different bacterial species such as *Streptomyces coelicolor* (Burrell *et al.,* 2012).

Polyamine biosynthesis depends on the activity of basic amino acid decarboxylases using Lysine, Arginine or Ornithine as specific substrate to produce Cad, Agmatine (Agm) or Put, respectively. These enzymes can therefore be generally referred to as LAOdcs (Lysine-Arginine-Ornithine decarboxylases). In most eukaryotes, Put synthesis is carried out by an ornithine decarboxylase (Odc) (Michael, 2016a, Michael, 2016b). An alternative pathway for Put biosynthesis found in bacteria and plants involves decarboxylation of arginine by arginine decarboxylases (Adc); these enzymes produce Agm which is further converted into Put by agmatine iminohydrolase/deiminase and N-carbamoylputrescine amidohydrolase (Michael, 2016a, Michael, 2016b). Interestingly, plants (Lee & Cho, 2001, Bunsupa *et al.,* 2012) and some bacteria exemplified by S. *ruminantium* and *V. vulnificus* (Takatsuka *et al.,* 2000, Lee *et al.,* 2007) possess a bifunctional Odc/Ldc capable of synthesizing both Put and Cad. Spd is formed by spermidine synthase (SpdSyn) through aminopropylation of Put, using an aminopropyl group released by decarboxylation of S-adenosylmethionine (Michael, 2016a, Michael, 2016b). In addition, *E. coli* SpdSyn transforms Cad into aminopropyl-Cad, a Spd analogue sharing its growth stimulating properties (Kim *et al.,* 2016).

Two major structural protein super-families are responsible for polyamine biosynthesis through pyridoxal-5-phosphate (PLP)-dependent LAOdcs: the alanine racemase fold (AR-fold) super-family and the aspartate aminotransferase fold (AAT-fold) super-family (Eliot & Kirsch, 2004). Phylogenetic studies of the LAOdcs of the AR-fold super-family revealed that these enzymes are widespread throughout the three domains of life while the AAT-fold LAOdcs are found exclusively in Bacteria and a few archaea (Lee *et al.,* 2007, Burrell *et al.,* 2010). Bacterial AAT-fold LAOdcs can be divided in two types according to the presence or absence of a particular CheY-like response regulator receiver domain, known as the “wing domain”, necessary for the formation of higher-order oligomers (Burrell *et al.,* 2010, Kanjee *et al.,* 2011b). The short form referred as wing-less LAOdc was found in *Firmicutes, Cyanobacteria* and *Actinobacteria* phyla, and a few “wing-less” AAT-fold decarboxylases from *Firmicutes* and *Actinobacteria* were shown to have an Adc activity, required in particular for biofilm formation in *Bacillus subtilis (Firmicutes)* (Burrell *et al.,* 2010). The long, wing domain-containing form likely originated in *Proteobacteria* (Kanjee *et al.,* 2011b).

AAT-fold decarboxylases with the wing domain have been intensively studied in *Enterobacteria* since the early 1940s (Gale & Van Heyningen, 1942, Gale & Epps, 1944) because of the link between enterobacterial pathogenicity for humans and their capacity to withstand acid stress thanks to the crucial role of the inducible LAOdcs. Consequently, the current understanding of the AAT-fold LAOdc is based on analyses of a very limited number of bacterial species, *i.e.* mostly *enterobacteria: E. coli, Salmonella typhimurium, Vibrio cholerae* and *V. vulnificus.* At a specific acidic pH, expression of the decarboxylase gene is induced by an excess of the target amino acid uptaken by a dedicated inner membrane antiporter (Kanjee & Houry, 2013). The enzyme transforms the amino acid substrate into the corresponding polyamine upon consumption of a proton and production of a CO_2_ molecule. In association with the polyamine excretion by the antiporter, this reaction results in an efficient buffering of the intracellular medium and the extracellular surroundings. These inducible stress response decarboxylases are distinguished from “biosynthetic” enzymes that are involved only in polyamine biosynthesis. *E. coli* encodes two biosynthetic decarboxylases (LdcC: Lys->Cad and OdcC/SpeC: Orn->Put) responsible for Cad and Put biosynthesis respectively, and three acid stress-inducible decarboxylases (Ldcl: Lys->Cad, AdcI/AdiA: Arg->Agm and Odcl/SpeF: Orn->Put) which together constitute a very robust acid stress response system that allows the bacterium to survive upon acid stress as low as pH 2.0 (Zhao & Houry, 2010, Kanjee *et al.,* 2011a, Kanjee & Houry, 2013). The importance of LdcI was also demonstrated in *S. typhimurium, V. cholerae* and *V. vulnificus,* where it promotes growth and survival under acidic conditions but also confers protection from oxidative stress insults (Merrell & Camilli, 2000, Kang *et al.,* 2007, Viala *et al.,* 2011)). It should be noted that LdcI-encoding genes are often designated as *cadA* as originally proposed upon identification of this gene in *E. coli* (Tabor *et al.,* 1980).

*E. coli* LdcI but not LdcC interacts with the AAA+ ATPase RavA to assemble into a huge macromolecular cage (Snider *et al.,* 2006, El Bakkouri *et al.,* 2010, Malet *et al.,* 2014, Kandiah *et al.,* 2016). One of the functions of this mysterious complex is to protect LdcI from inhibition by the stringent response alarmone ppGpp, thus enabling the bacterium to efficiently cope with both acid and nutrient stress simultaneously (El Bakkouri *et al.,* 2010, Kanjee *et al.,* 2011a, Malet *et al.,* 2014). While investigating why RavA binds only LdcI but not LdcC, we documented numerous inconsistencies in annotation of enterobacterial *ldcl* and *IdcC* genes and realized that each of these two families appeared to have a distinct genetic context (Kandiah *et al.,* 2016). Thus, spurred on by the limited nature of the previous studies, we set out to perform an extensive phylogenetic analysis of the AAT-fold LAOdcs in circa 4,500 complete prokaryotic proteomes to decipher the evolutionary history of these proteins and their functional evolution. In the present study, we revealed the ancestral nature of the wing-less LAOdcs in *Cyanobacteria* and *Firmicutes,* and the complex evolutionary history of long AAT-fold LAOdcs in *Proteobacteria,* leading to the emergence of distinct subfamilies. Moreover, we disclosed a novel subfamily of enzymes, clearly distinct from the well-known LdcI, LdcC, Adcl, Odcl and OdcC families, but more related to Ldc and Adc than to Odc. Excitingly, this novel evolutionary subfamily has been overlooked in previous phylogenetic analyses in spite of its wide distribution in *Betaproteobacteria* and *Pseudomonadaceae,* implying that it deserves a thorough characterization and a functional comparison with the known AAT-fold long LAOdc. The only previously mentioned LAOdc from these taxa is the lysine decarboxylase LdcA from a major multidrug resistant opportunistic human pathogen *Pseudomonas aeruginosa* (Chou *et al.,* 2010). Thus, here we went beyond the initial characterization of the *P. aeruginosa* LdcA (Chou *et al.,* 2010), and further analyzed its expression, regulation, and function in the light of the available knowledge in particular on *E. coli* LdcI and LdcC. This combined phylogenetic and functional study revealed that LdcA belongs to a novel subgroup of the long AAT-fold LAOdcs, and that its function is linked to Cad production and to the general polyamine metabolism rather than to stress response.

## Results

### Taxonomic distribution of AAT-fold LAOdcs in prokaryotes

An in-depth survey of 4,466 prokaryotic proteomes representing 1,904 species revealed 4,090 protein sequences belonging to the AAT-fold LAOdcs (13 of which were unannotated or annotated as pseudo-genes) (Fig. S1A). Representatives of this super-family are mainly present in Bacteria, especially in *Proteobacteria, Firmicutes, Actinobacteria* and *Cyanobacteria,* while only seven sequences were detected in *Archaea* (Fig. S1A), indicating that AAT-fold LAOdc are very likely of bacterial origin. The corresponding Maximum Likelihood (ML) tree could be divided in three parts (Fig. S1B). Cluster I encompasses sequences devoid of the wing domain (short AAT-fold LAOdc) (Bootstrap value (BV) = 99%). They are mainly found in *Firmicutes, Cyanobacteria* and *Actinobacteria.* Cluster II gathers nearly all proteobacterial sequences and a few sequences from *Firmicutes* (BV = 99%), all containing a wing domain. Cluster III branches in-between Cluster I and Cluster II; it is composed of a mix of long and short AAT-fold LAOdc sequences from unrelated taxonomic groups *(Proteobacteria, Actinobacteria, Bacteroidetes,* and *Firmicutes),* strongly suggesting that they spread through horizontal gene transfers among these lineages.

### Wingless AAT-fold LAOdc are ancestral in Firmicutes and Cyanobacteria

Within Cluster I, all cyanobacterial sequences group together albeit with a weak bootstrap value (BV) (Fig. 1A). They are widely distributed in this phylum (Fig. 2). The phylogeny inferred with these sequences (Fig. 2A) is overall consistent with a reference phylogeny of *Cyanobacteria* based on ribosomal proteins (Fig. 2B). This indicates that a gene coding for a wing-less AAT-fold LAOdc was likely present in the ancestor of *Cyanobacteria* and had been mainly transmitted vertically in *Cyanobacteria.* The genomic context of LAOdc in *Cyanobacteria* is not conserved even in relatively close species (Fig. 2C).

**FIG 1.**
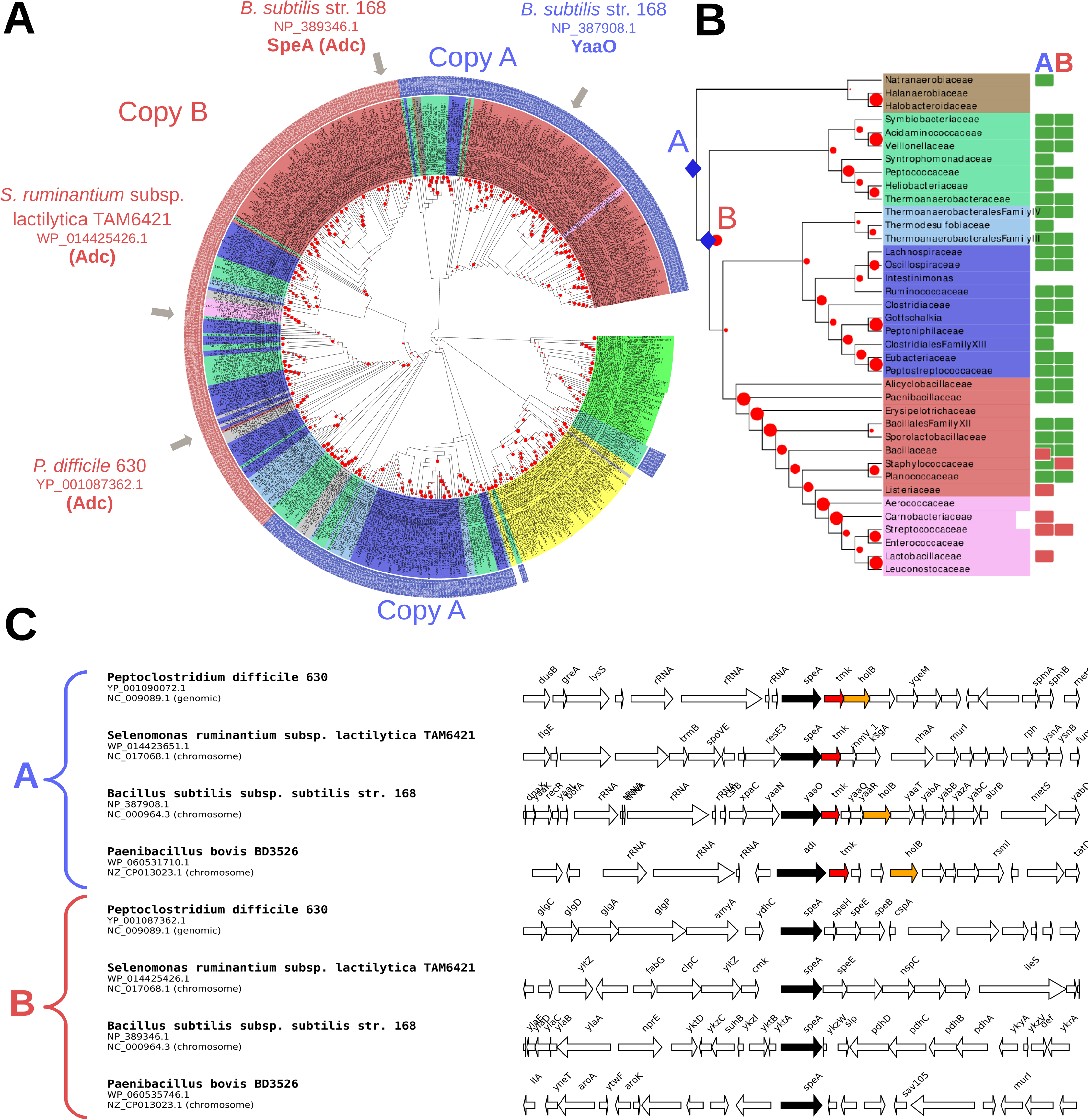
LAOdcs are ancestral in *Firmicutes.* (A) Unrooted maximum likelihood phylogeny of LAOdc Cluster I (PhyML, LG+I+G4, 504 sequences, 295 amino-acid positions), displayed as a cladogram. The corresponding phylogram is available at the newick format as supplementary data. Leave colors correspond to taxonomic groups *(Firmicutes:* red, *Cyanobacteria:* green, *Actinobacteria:* yellow, other: grey). External colored rings correspond to copy A (blue) and B (pink). LAOdc sequences discussed in the text or for which functional information is available are indicated with gray arrows. Red dots correspond to bootstrap values (BV). The size of the dots is proportional to BV. (B) Taxonomic distribution of AAT-fold LAOdc according to a reference maximum likelihood phylogeny of *Firmicutes,* displayed as a cladogram. The corresponding phylogram is available at the newick format as supplementary data. The reference tree was inferred using ribosomal protein sequences (PhyML, LG+I+G4, 38 sequences, 6,133 amino-acid positions) and rooted according to a recent study by Antunes *et al.* (Antunes *et al.,* 2016) Red dots correspond to bootstrap values (BV). The size of the dots is proportional to BV. The blue diamonds pinpoint the emergence of copy A and copy B. Rectangles at leaves indicate that at least one genome of the considered taxon encodes one or more AAT-fold LAOdc. More precisely a green rectangle indicates that the ancestor of the taxon likely contains one (or more) AAT-fold LAOdc gene, while a red rectangle indicates that some members of the taxon acquired secondarily their AAT-fold LAOdc by horizontal gene transfer. (C) Genomic context of LAOdc A and B in a sample of *Firmicutes.* Black arrows: LAOdc coding genes, colored arrows: conserved neighbor genes.

**FIG 2.**
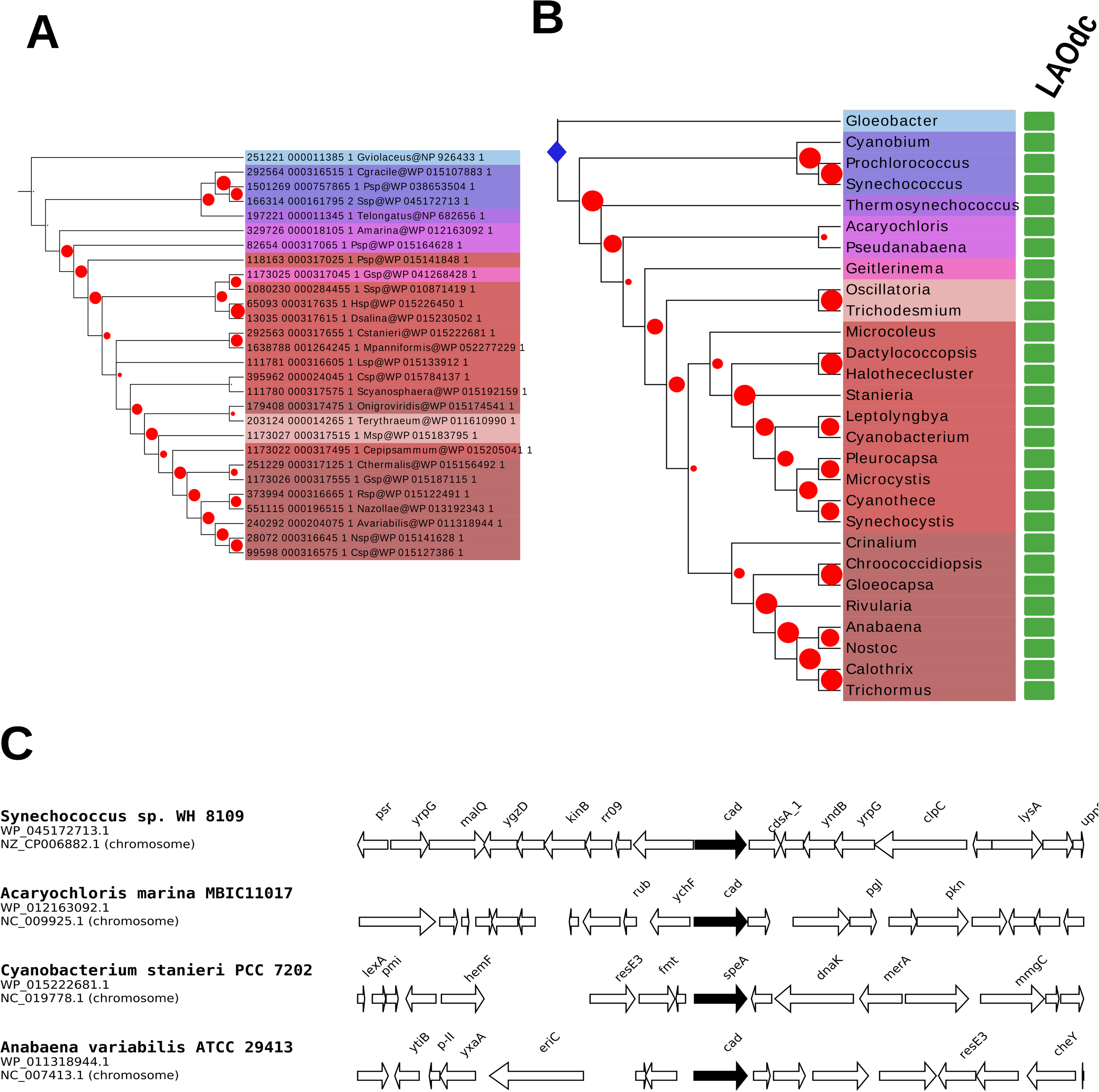
LAOdcs are ancestral in *Cyanobacteria.* (A) Maximum likelihood phylogeny of LAOdc of *Cyanobacteria* (PhyML, LG+I+G4, 28 sequences, 446 amino-acid positions), displayed as a cladogram. The corresponding phylogram is available at the newick format as supplementary data. The tree has been rooted according to the reference phylogeny of *Cyanobacteria* (see above). Leave colors correspond to taxonomic groups. Red dots correspond to bootstrap values (BV). The size of the dots is proportional to BV. (B) Taxonomic distribution of AAT-fold LAOdc according to a reference maximum likelihood phylogeny of *Cyanobacteria,* displayed as a cladogram. The corresponding phylogram is available at the newick format as supplementary data. The reference tree was inferred using ribosomal protein sequences (PhyML, LG+I+G4, 30 sequences, 6,394 amino acid positions). The tree has been rooted using sequences from *Firmicutes (Natranaerobius thermophilus)* and *Actinobacteria (Streptomyces albulus).* Red dots correspond to bootstrap values (BV). The size of the dots is proportional to BV. The blue diamond pinpoints the origin of AAT-fold LAOdc in *Cyanobacteria.* Rectangles at leaves indicate that at least one genome of the considered taxon encodes one or more AAT-fold LAOdc. More precisely a green rectangle indicates that the ancestor of the taxon likely contains one (or more) AAT-fold LAOdc gene. (C) Genomic context of LAOdc in a sample of *Cyanobacteria.* Black arrows: LAOdc genes.

Similarly, sequences of *Firmicutes* belonging to Cluster I are widely distributed in this phylum (Fig. 1). The comparison of the phylogeny inferred with these sequences (Fig. 1A) with a reference phylogeny of *Firmicutes* (Fig. IB) suggests that a gene coding for a short AAT-fold LAOdc was present in the ancestor of this phylum. Noteworthy, most of members of *Firmicutes* harbor two LAOdc copies (referred as A and B), with exception of the most early-branching lineages: one single gene corresponding to the copy A is found in *Natranaerobiaceae,* while no gene is found in *Halobacteroidaceae* and *Halanaerobiaceae* (Fig. IB). These two copies can be easily distinguished by their genomic context (Fig. 1C): the copy A presents a well-conserved association with thymidylate kinase and DNA polymerase III subunit delta coding genes, while the genomic context of copy B was not conserved. The presence of gene coding for a copy A in the early diverging *Natranaerobiaceae* lineage and in most *Firmicutes* taxa suggests that the copy A could be ancestral in *Firmicutes.* In contrast, the copy B seems to appear in the common ancestor shared by *Clostridia* and *Bacilli,* possibly as the result of a gene duplication event. Worthy of note, a loss of both copies can be inferred in the ancestor of *Lactobacillales.* Yet, some *Streptococcaceae* and *Carnobacteriaceae* have reacquired either copy A or copy B by horizontal gene transfer from different *Firmicutes* donors (Fig. 1A). Finally, Cluster I encompassed a group of sequences from *Actinobacteria.* Their taxonomic distribution is patchy and their phylogeny is not consistent with the current taxonomy, suggesting that they have been acquired and spread through horizontal gene transfer in *Actinobacteria.*

### Wing-domain containing AAT-fold LAOdc form four groups in Proteobacteria

The phylogenetic analysis of proteobacterial AAT-fold LAOdcs composing the Cluster II revealed two monophyletic groups, corresponding to Ode (Posterior probabilities (PP) = 1, Fig. 3A and BV = 100%, Fig. 3B) and LAdc (PP = 1, Fig. 3A and BV = 100%, Fig. 3B). The LAdc group is further split into Ldcl/LdcC (PP = 1, Fig. 3A and BV = 100%, Fig. 3B), Adc (PP = 1, Fig. 3A and BV = 93, Fig. 3B), and a hitherto non-identified subfamily (PP = 1, Fig. 3A and BV =100%, Fig. 3B), that will be referred as LdcA in the name of its only previously mentioned member, LdcA from *P. aeruginosa* (Chou *et al.,* 2010). It should be emphasized that although the monophyly of Odcl/OdcC, Ldcl/LdcC, Adc and LdcA groups is well supported, the relationships among Ldcl/LdcC, Adc and LdcA are unresolved (PP = 0.67, Fig. 3A and BV < 80%, Fig. 3B).

**Fig. 3.**
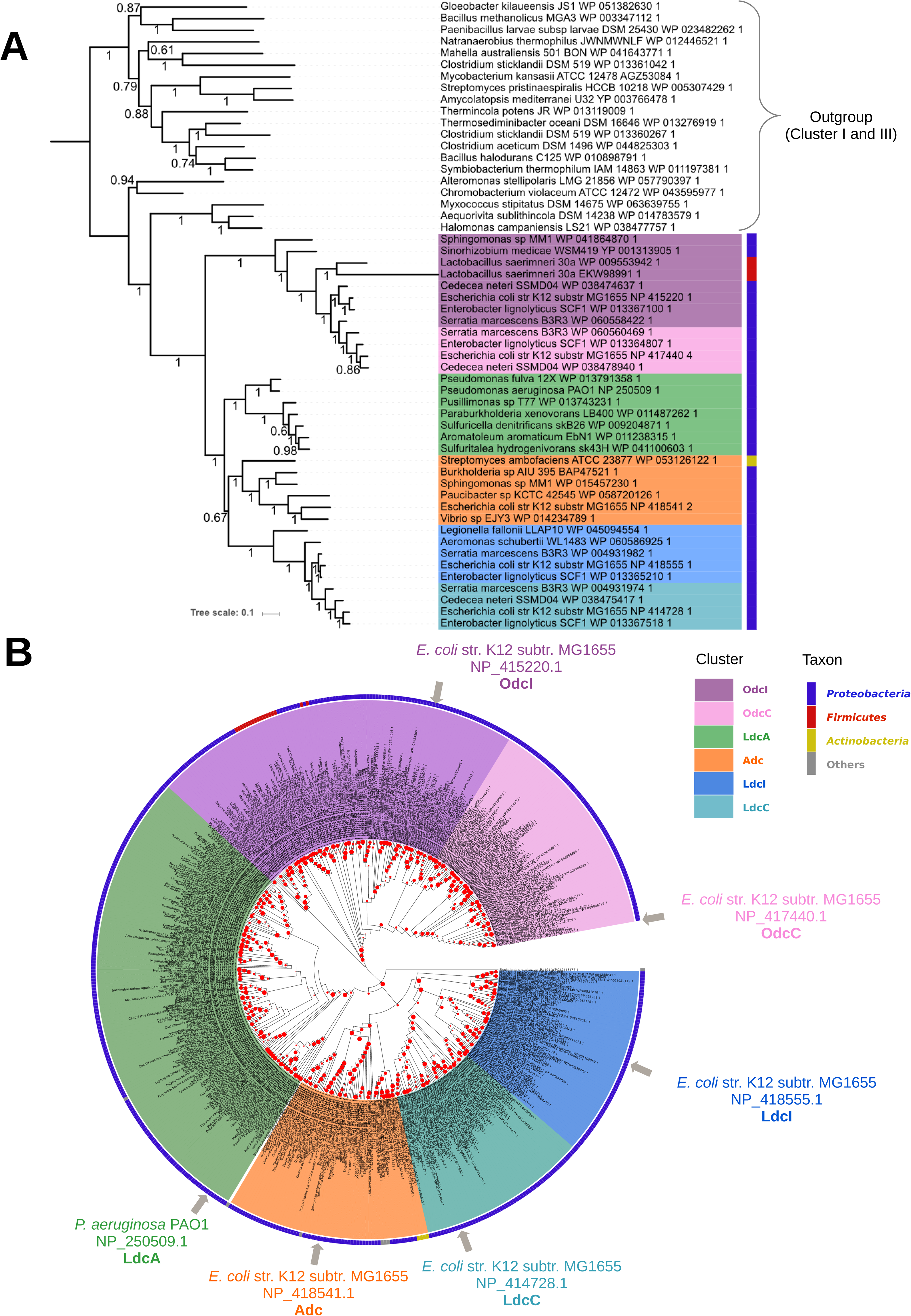
Phylogeny of the LAOdc Cluster II (A) Bayesian phylogeny of cluster II inferred from a sample of representative sequences and rooted with a sample of sequences from clusters I and III (MrBayes, mixed model+G4, 54 sequences, 392 amino acid positions). The scale bar represents the average number of substitutions per site. Numbers at branches correspond to posterior probabilities. The ML tree inferred with the same dataset supported the same topology (see supplementary data). (B) Maximum likelihood phylogeny of the LAOdc Cluster II (PhyML, LG+I+G4, 551 sequences, 589 amino-acid positions), displayed as a cladogram. The corresponding phylogram is available at the newick format as supplementary data. LAOdc sequences from *E. coli* and *P. aeruginosa* discussed in the text are indicated with gray arrows. The tree has been rooted according to (A). Colors on the external circle correspond to taxonomic groups: dark blue: *Proteobacteria,* red: *Firmicutes,* yellow: *Actinobacteria,* gray: other taxa. Red dots correspond to bootstrap values (BV). The size of the dots is proportional to BV.

The comparison of the LAOdc phylogeny (Fig. 3B) and taxonomic distribution with a reference phylogeny of *Proteobacteria* (Fig. 4A) suggests that *odd* could be ancestral in *Bradyrhizobiaceae, Xanthobacteriaceae, Methylobacteriaceae,* and *Beijerinckiaceae* (all belonging to *Alpha proteobacteria),* as well as in *Vibrionaceae, Pasteurellaceae,* and *Enterobacteriaceae* (all belonging to *Gammaproteobacteria).* In contrast, *odcC* seems more recent and appears to result from a gene duplication of *odd* that occurred in *Enterobacteriaceae,* before the divergence of *Sodalis* (PP = 1, Fig. 3A and BV = 75%, Fig. 3B, and Fig. 4B). Genes coding for Odd and OdcC are clearly distinguished by their context (Fig. S2). More precisely, *odd* is predominantly upstream from a gene *(potE)* coding for a putrescine-ornithine antiporter, consistently with the function of Odd that converts Orn to Put. In contrast, OdcC coding genes are located in the vicinity of Fe^2^+-trafficking protein *(yggX),* a lytic murein transglycosylase *(mltC)* genes, an A/G-specific adenine glycosylase *(mutY),* a tRNA (guanosine(46)-N7)-methyltransferase *(trmB)* and an hypothetical protein. Interestingly, a few horizontal gene transfers of *odd* occurred from *Proteobacteria* to unrelated *firmicutes:* some *Lactobacillus, Staphylococcus lugdunensis,* and *Megasphaera elsdenii.*

**Fig. 4.**
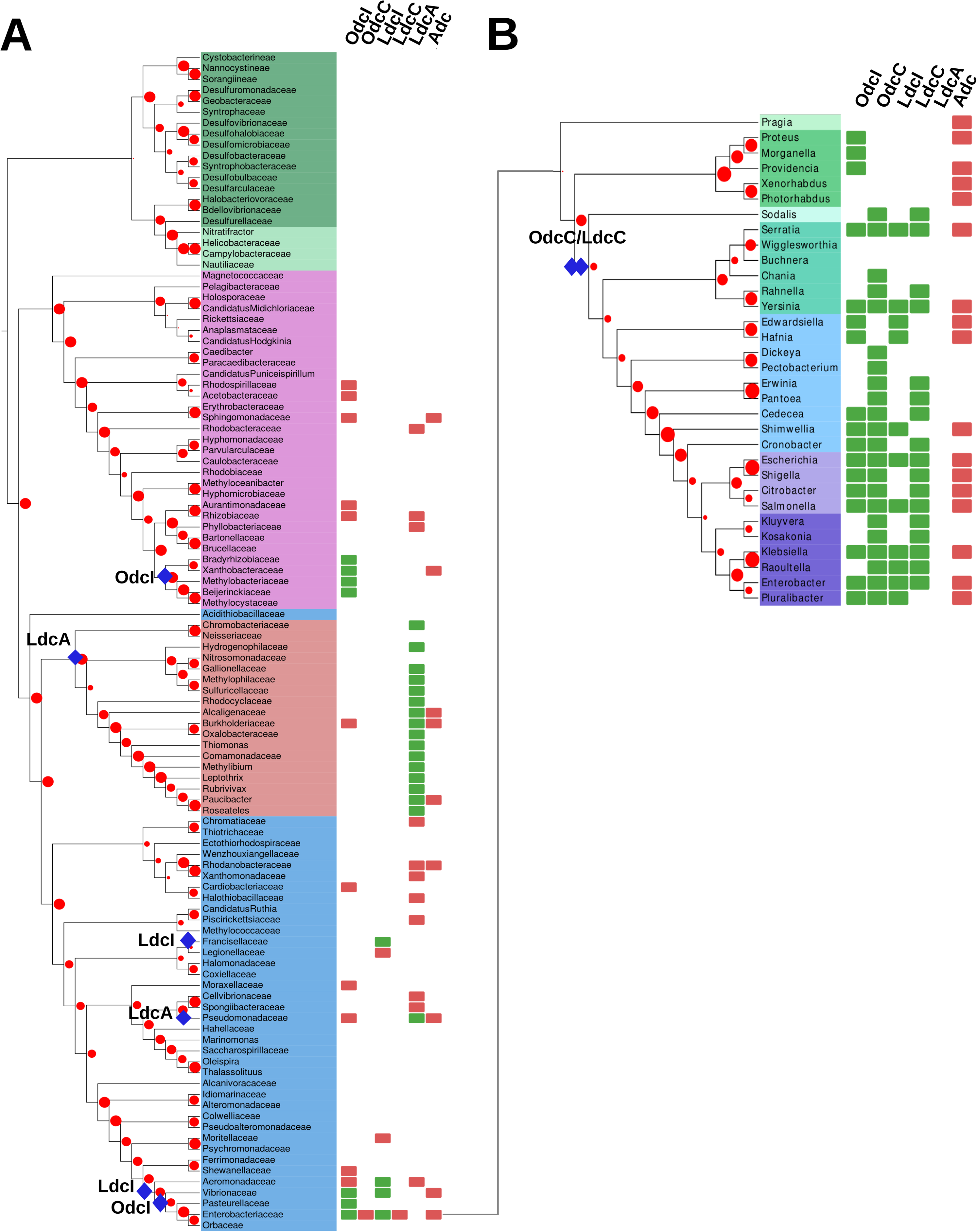
Taxonomic distribution of LAOdcs in *Proteobacteria* (A) Taxonomic distribution of AAT-fold LAOdc according to a reference maximum likelihood phylogeny of *Proteobacteria,* displayed as a cladogram. The corresponding phylogram is available at the newick format as supplementary data. The reference tree was inferred using ribosomal protein sequences (PhyML, LG+I+G4, 108 sequences, 6,129 amino acid positions). The tree has been rooted in the branch separating *Deltaproteobacteria* and *Epsilonproteobacteria* in agreement with the study by Gupta (Gupta, 2000). Leave colors correspond to taxonomic groups. Red dots correspond to bootstrap values (BV). The size of the dots is proportional to BV. A blue diamond indicates the ancestral presence of LAOdc families in the corresponding taxon. Rectangles at leaves indicate that at least one genome of the considered taxon encodes one or more AAT-fold LAOdc. More precisely a green rectangle indicates that the ancestor of the taxon likely contains one (or more) AAT-fold LAOdc gene, while a red rectangle indicates that some members of the taxon acquired secondarily their AAT-fold LAOdc by horizontal gene transfer. (B) Taxonomic distribution of AAT-fold LAOdc according to a reference maximum likelihood phylogeny of *Enterobacteraceae,* displayed as a cladogram. The corresponding phylogram is available at the newick format as supplementary data. The reference tree was inferred using ribosomal protein sequences (PhyML, LG+I+G4, 34 sequences, 6,333 amino acid positions). The tree was rooted with *Shewanella baltica (Alteromonadales)* and *Pasteurella multocida (Pasteurellales).* Leave colors correspond to taxonomic groups. Other legend elements are identical to (A).

Ldcl, also called *cadA,* appears to be ancestral in some gammaproteobacterial lineages (*Francisellaceae, Aeromonadaceae, Vibrionaceae* and *Enterobacteriaceae,* Fig. 4A). Moreover, similar to *odd/odcC, ldcC* apparently derived from *ldcl,* and more precisely from a gene duplication that occurred, as for *odd,* just before the emergence of *Sodalis* (PP = 1, Fig. 3A and BV = 100%, Fig. 3B, and Fig. 4B). The *Idcl* gene forms the *cadBA* operon together with the lysine-cadaverine antiporter-encoding the *cadB* gene. In *Enterobacteriaceae,* this operon is known to be regulated by the transcriptional factor CadC integrating three external signals - low pH, high lysine and low Cad levels (Kuper & Jung, 2005, Fritz *et al.,* 2009). Our analysis confirmed the conserved genomic organization of the *cadCBA* system (Fig. S2) (Zhao & Houry, 2010) allowing *Enterobacteria* to face acid and oxidative stresses. The *ldcC* genomic context seems to be also strongly conserved (Fig. S2) with genes coding for hydroxymyristol acyl carrier protein dehydratase, UDP-N-acetylglucosamine acetyltransferase, tetraacyldisaccharide-1-P synthase, ribonuclease Hll, DNA polymerase III alpha subunit, acetyl-CoA carboxylase 2C alpha subunit upstream and putative lyase and tRNA(lle)-lysidine synthetase genes downstream.

In sharp contrast with OdcI/OdcC and LdcI/LdcC, the Adc group presents a patchy taxonomic distribution and the relationships among Adc sequences are at odds with current systematics, with sequences from different classes of *Proteobacteria* being intermixed on the tree (Fig. 3B and Fig. 4). This indicates that the Adc subfamily was heavily impacted by horizontal gene transfers. The genomic context of *adc* is not conserved (Fig. S2), precluding identification of potential functional partners. Interestingly, the *Burkholderia* sp. AIU 395 LAOdc that locates in the Adc group was shown to possess a lysine decarboxylase and oxidase activity (Sugawara *et al.,* 2014).

### Phylogenetic analyses of LAOdc disclose a fourth group in Proteobacteria

As specified above, beside OdcI/C, LdcI/C and Adc, a fourth group of LAOdc sequences is present in the tree. This group contains LdcA from *P. aeruginosa,* shown to possess a lysine decarboxylase activity (Chou *et al.,* 2010). Homologues of LdcA are found in other *Pseudomonadaceae* and in *Betaproteobacteria.* Both groups of sequences are well separated on the tree and widely distributed in these two taxa, with relationships globally in agreement with the current taxonomy. This suggests that an LdcA homologue was present in the ancestor of *Pseudomonadaceae* and in the ancestor of *Betaproteobacteria* and has been globally well conserved along the diversification of these two taxa. The genetic environment of *IdcA* is not conserved (Fig. S2). To our knowledge, the only publication describing a member of the LdcA subfamily concerns *P. aeruginosa,* a highly versatile bacterium that efficiently grows on arginine but not on lysine (Fothergill & Guest, 1977, Rahman & Clarke, 1980). In this organism exhibiting tightly interconnected lysine and arginine catabolism networks (Chou *et al.,* 2010, Madhuri Indurthi *et al.,* 2016), the *PA1818* gene was identified as a part of the ArgR regulon upon growth on excess arginine, but was surprisingly found to code for a lysine decarboxylase, and not for an arginine decarboxylase as one would have logically supposed, and therefore called *ldcA* (Chou *et al.,* 2010, Madhuri Indurthi *et al.,* 2016). *P. aeruginosa* is a major cause infections, especially in patients with compromised and weakened immune system. This opportunistic pathogen is also a well-identified threat for patients suffering from Cystic Fibrosis (CF), because the chronic respiratory infections associated to host inflammatory responses lead to pulmonary tissue destruction and lung failure (Bodey *et al.,* 1983, Gellatly & Hancock, 2013). The occurrence and persistence of *P. aeruginosa* in the CF patients’ lungs, whose secretions were shown to be acidified and to become oxidative (Pezzulo *et al.,* 2012), hints to a possible role of LdcA in promoting bacterial fitness. Therefore, in the following sections, we chose to deepen the present knowledge on expression, regulation and functional characterization of *P. aeruginosa* LdcA.

### LdcA expression is growth-phase dependent

To determine the role of the LdcA protein in *P. aeruginosa,* the prerequisite was to know when the protein is produced by the bacterium and what are the mechanisms controlling its production. Thus, we first analyzed the expression of its gene by creating a transcriptional fusion between the *IdcA* promoter (*PldcA)* and the reporter *lacZ* gene. The fusion was then integrated into the chromosome of PAO1 (see Materials and Methods for further details), the first *P. aeruginosa* sequenced strain (Stover *et al.,* 2000) considered as a reference strain and widely studied. We then measured the β-galactosidase activity of the PAO1::*P*_*idcA*_*-lacZ* strain grown in a minimal medium P (MMP) containing glutamate as a carbon source and lysine or arginine (20mM) as additives. Unlike lysine, arginine was able to induce the expression of *IdcA* in PAOl:*:P*_*idcA*_*-lacZ* (not shown), in agreement with published data identifying ArgR as a positive regulator of *IdcA* expression in the PAO1 strain (Lu *et al.,* 2004, Chou *et al.,* 2010).

The pattern of *ldcA* expression was followed during the growth both in minimal and rich media. In MMP medium supplemented with glutamate and arginine (MMP-GR), the β-galactosidase activity increased slightly but continuously along the growth and reached a maximum in the stationary phase (Fig. 5A). Expression of *ldcA* in LB rich medium followed the same pattern of expression (Fig. 5B), that was paralleled by an increase in LdcA protein amount assessed by western blot (not shown). The expression level obtained in LB was two-fold lower that the one measured in MMP-GR. Addition of 20 mM arginine to LB did not change the growth rate of the bacteria but the *ldcA* promoter activity increased during the transition to the stationary phase to reach a level similar to the one measured in MMP-GR (Fig. 5B), indicating a probable limiting concentration of the amino acid in LB at late growth.

**Fig. 5.**
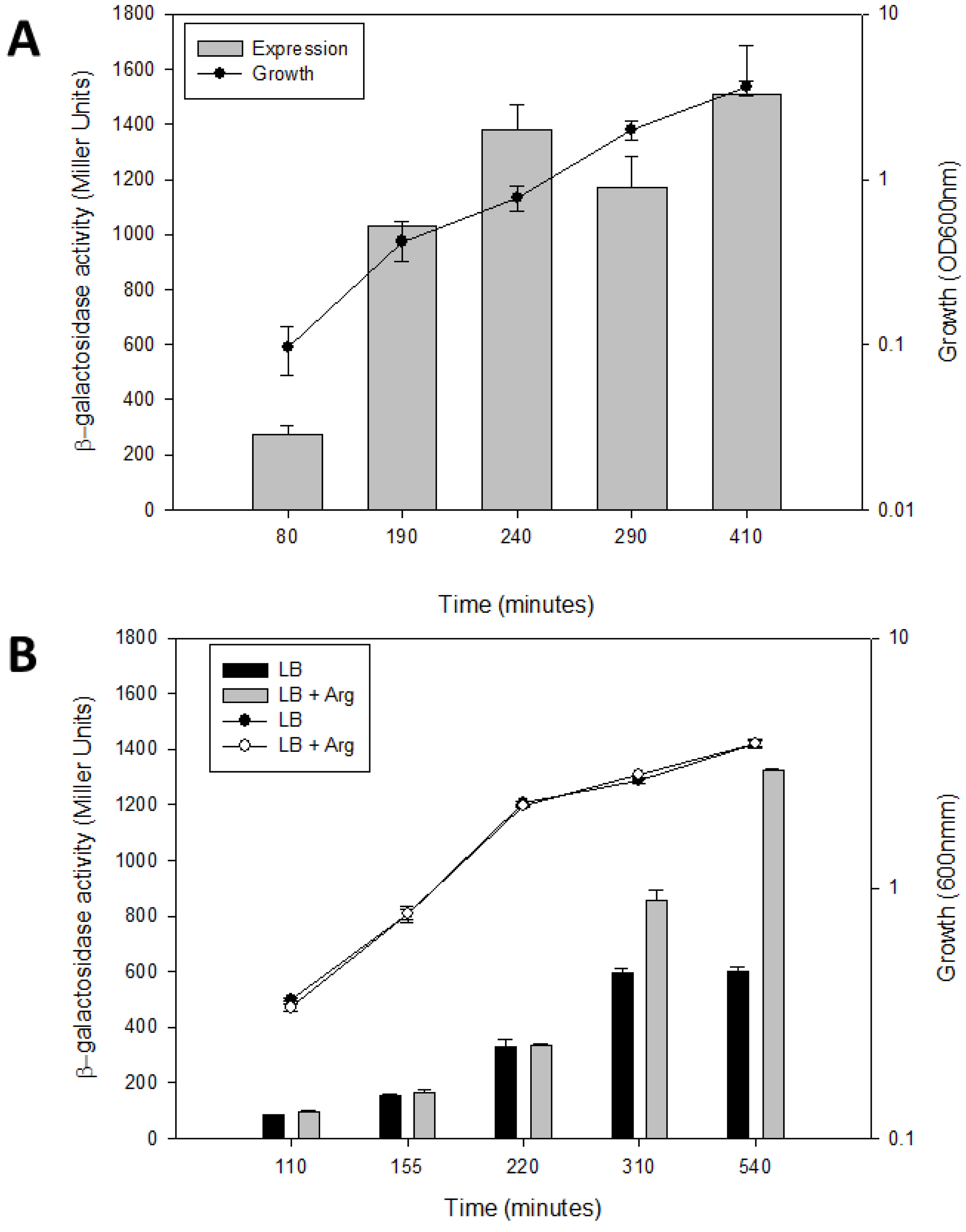
Factors influencing *IdcA* expression Activity of *IdcA* promoter fused to *lacZ* reporter gene was assessed during growth in different media and genetic backgrounds. Measurements of theβ-galactosidase activity of *PA01::PldcA-lacZ* strain grown either in minimal medium P (MMP) containing 20 mM L-glutamate and 20 mM arginine (A), or in LB containing or not 20 mM arginine (B) were performed at times indicated. C.β-galactosidase activity of wild-type and indicated mutant CHA strains harboring the *PldcA-lacZ* fusion and grown in LB.β-galactosidase activity is expressed in Miller Units (left Y-axis) and presented in the bar graphs. Growth was performed in 125 ml flasks, followed by measure of OD_600_ (right Y-axis) and plotted on lines. Results are the average of values from three independent experiments ⍰±⍰ standard deviation (SD).

### LdcA expression differs from that of LdcC and Ldcl

The *IdcA* genomic context being different from that of *ldcl* and *IdcC* genes (Fig. S2), we wondered if *ldcA* was also regulated differently than these genes; this knowledge could further provide information on its role in *P. aeruginosa* physiology. Even if *E. coli* LdcC is called “constitutive Lysine Decarboxylase”, the corresponding gene was shown to be induced in stationary phase in LB medium by RpoS, the sigma factor of stationary phase (Kikuchi *et al.,* 1998). However, *ldcA* expression was not affected in a *rpoS* mutant background (data not shown), in agreement with previous transcriptomic analyses demonstrating that *ldcA* is not part of the RpoS regulon (Schuster *et al.,* 2004). To compare with LdcI, the “inducible Lysine Decarboxylase”, despite the absence of a CadC homologue in *P. aeruginosa,* we assessed if acid stress could activate the expression of *ldcA* in a CadC-independent manner. Hence, we induced acid stress by decreasing the pH of the medium to a value of 5 during the exponential phase of growth and documented an absence of effect of this treatment on *ldcA* expression (Fig. S3A). In addition, *ldcI* from *V. vulnificus* was reported to be induced by SoxR upon H_2_0_2_ stress (Kim *et al.,* 2006). In *P. aeruginosa,* SoxR is not a key player in the oxidative stress response, but H_2_0_2_ activates the global regulator OxyR that orchestrates the defense against ROS (Ochsner *et al.,* 2000) conclude, *P. aeruginosa* does not overexpress *ldcA* to respond to low pH or oxidative stress conditions.

### Low pH survival is not affected in absence of LdcA

To clarify the function of LdcA in *P. aeruginosa,* a mutant deleted of the *ldcA* gene, as well as a complemented strain in which one copy of *ldcA* driven by its own promoter was inserted in the chromosome, were engineered (see Materials and Methods). Using Biolog system, a Phenotype MicroArray analysis was carried out to test a large number of stress conditions in systematical and reproducible manner. A potential role of LdcA in detoxifying and protecting *P. aeruginosa* during acid, alkaline and oxidative stress, antibiotic treatment and toxic molecules causing DNA damage, nitrosative stress, and membrane destabilization was assessed. After monitoring the growth of *P. aeruginosa* in the different conditions in minimal medium during 24 hours (see Materials and Methods), the “Area Under the Curve” (AUC) in each condition was calculated and compared between the strains. Analysis of the growth fitness of wild-type, mutant and complemented strains indicated that *P. aeruginosa* could grow optimally in a pH range from 5 to 10 without considerable effect on the metabolism and biomass growth. At pH 4 to 5, the bacterial growth started to be strongly inhibited and the strains were unable to grow below pH 4 (Fig. S3B). This set of experiments did not show any significant difference between the mutant and the wild-type and complemented strains, pointing to a non-involvement of LdcA in survival at low pH. Similarly, no significant effect of *ldcA* absence on resistance against antibiotics, oxidative and toxic agents could be detected (not shown). Hence LdcA seems not to be important for stress response.

### The Cad pool generated by LdcA impacts persistence phenotype and polyamine content

In our quest for deciphering the importance of LdcA in bacterial physiology, we analyzed the role of its product Cad shown to play a fundamental role in *P. aeruginosa* virulence. Indeed, this polyamine was reported to be involved in the persistence of the bacterium which is of importance for its eradication by antibiotics; specifically, Cad production was shown to lead to a reduction of the dormant cells that form an antibiotic-tolerant subpopulation in MH medium (Manuel *et al.,* 2010).

Therefore, to assess the role of LdcA in persistence, we first confirmed the impact of *IdcA* mutation on Cad production in this rich MH medium. To do so, we quantified the intracellular Cad amounts in the bacterial strains by liquid chromatography coupled to high resolution mass spectrometry (LC/HRMS) during early-, mid-and late-exponential growth phases. Cad was completely absent in the *ldcA* mutant and complementation with a wild-type *ldcA* copy restored the metabolite level in the strain, indicating that, in PAO1, this polyamine is produced exclusively through the enzymatic activity of LdcA in this growth condition (Fig. 6). Moreover, the wild-type strain showed an impressive increase (around 25-fold) of Cad concentration during the growth, probably reflecting the *ldcA* expression pattern in rich medium (increase during exponential growth, Fig. 5B).

**Fig. 6.**
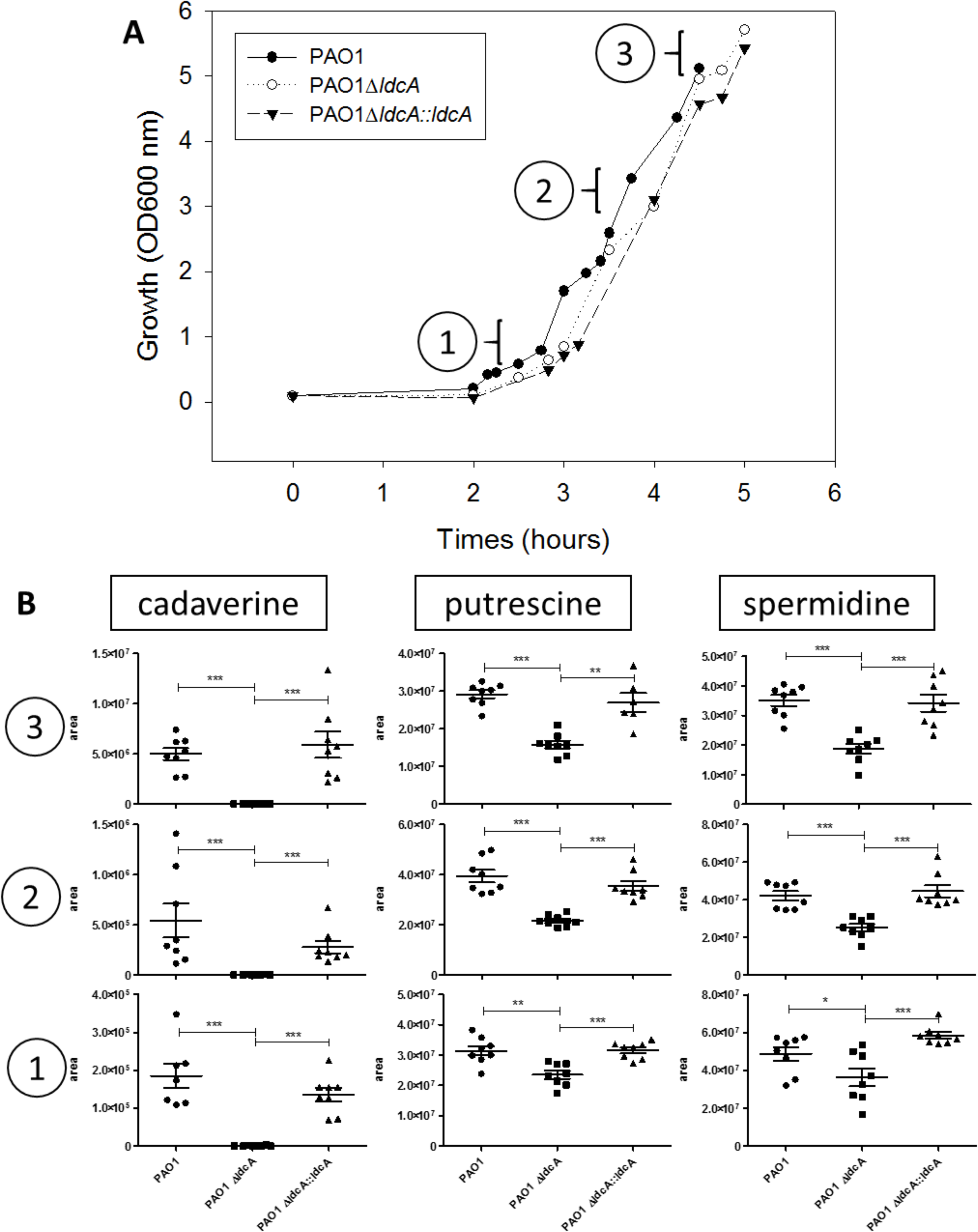
Intracellular Cad in rich medium is produced by the lysine decarboxylase LdcA A. Growth curves of the wild-type strain, the *ldcA* mutant and the complemented strain in the rich Mueller Hinton medium. B. Intracellular concentration, expressed in area of chromatogram peak, of the three indicated polyamines at (1) early-, (2) mid-and (3) late-exponential growth phases as indicated.

Then, the impact of *ldcA* on the number of persisters during carbenicillin treatment was assessed. As anticipated, *ldcA* mutant showed a number of persisters significantly higher compared to the wild-type PAO1 and complemented strains (Fig. S4), confirming the importance of LdcA activity in the persistence phenotype.

Interestingly, in parallel to Cad, we also quantified the intracellular concentrations of two other polyamines, Put and Spd, in the wild-type strain and *ldcA* mutant (Fig. 6). While in the wild-type PAO1, the amount of Cad is clearly growth-phase dependent, the amounts of Put and Spd were found to be abundant and constant in the MH medium. Remarkably, inactivation of *IdcA* affected not only the production of Cad but also the amount of intracellular Put and Spd which were reduced by around two-fold in the *ldcA* mutant at late exponential growth. This could either indicate a higher turnover of polyamine metabolism in the *ldcA* mutant or a reduced production of Put and Spd as a compensation for the absence of Cad.

### Polyamines are important for growth fitness

As mentioned above, LdcA synthesis is induced in minimal MMP medium in presence of arginine. Therefore, we monitored the growth of the *ldcA* mutant in this condition in presence of either lysine, the substrate of LdcA, or Cad, the product of its enzymatic transformation. The observed clear reduction of growth of the *ldcA* mutant highlighted the metabolic role of the enzyme, whereas wild-type growth was restored in the complemented strain (Fig. 7A). Addition of exogenous Cad was sufficient to restore a normal growth in the mutant indicating that the limiting factor was indeed the polyamine product (Fig. 7B). Strikingly, growth was also restored when Put (Fig. 7C) or Spd (Fig. 7D) were added to the growth medium at the same concentrations, indicating that these polyamines can substitute for Cad as a growth factor.

**Fig. 7.**
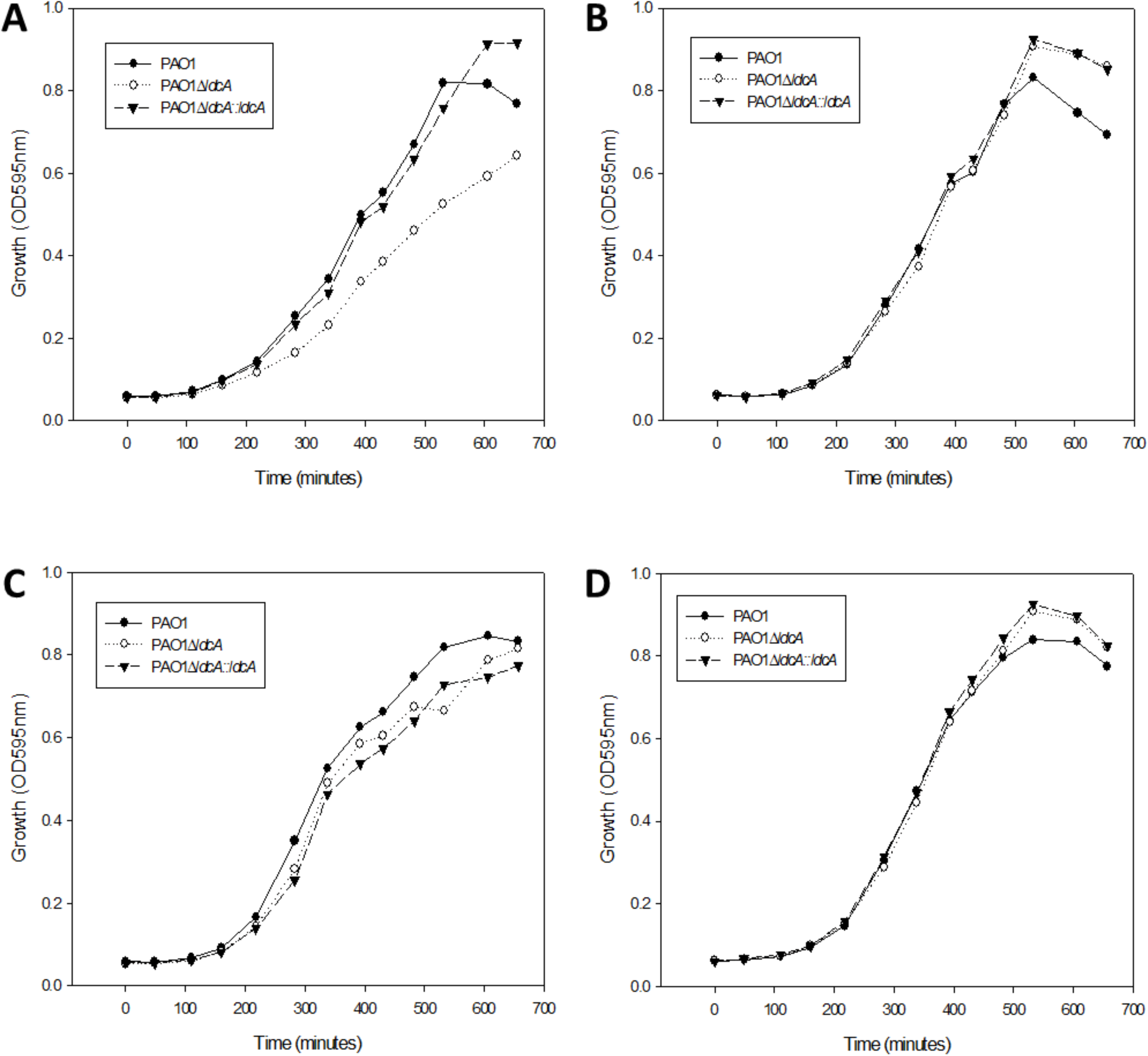
Polyamines are important for growth fitness in minimal medium Growth of the wild-type PAO1 strain, the *ldcA* mutant and the complemented strain in minimal MMP medium supplemented with 20 mM glutamate, ImM arginine and 5 mM of (A) lysine, (B) Cad, (C) Put or (D) Spd in 96-well plates. The experiments are representative of two experiments.

## Discussion

The present exhaustive phylogenetic analysis of the AAT-fold LAOdcs super-family indicates that wing-less enzymes were ancestral in *Firmicutes,* in agreement with earlier reports (Sekowska *et al.,* 1998, Burrell *et al.,* 2010). This analysis highlights also the ancestral presence of wing-less AAT-fold LAOdc in *Cyanobacteria.* Yet, no LAOdc activity of a cyanobacterial AAT-fold enzyme has been documented despite their purification from *Anabaena variabilis* and *Nostoc punctiforme* (Burrell *et al.,* 2010). We show that wing domain-containing AAT-fold LAOdc emerged during the diversification of *Proteobacteria,* suggesting that short AAT-fold LAOdc could be more ancient and that the acquisition of the CheY-like wing domain occurred likely secondarily, as previously proposed (Burrell *et al.,* 2010, Kanjee *et al.,* 2011a). The absence of AR-fold decarboxylases in *Firmicutes* (Burrell *et al.,* 2010, Michael, 2016a)implies that their AAT-fold decarboxylases would be the only route for polyamine biosynthesis and thus emphasizes the importance of these enzymes in this phylum. The *Firmicutes* AAT-fold LAOdc correspond to two copies, likely paralogues (A and B), present in most species and easily distinguishable by their genomic context. These two copies result from a duplication event that occurred very early during diversification of the phylum, probably in the ancestor of Clostridia and Bacilli (Fig. SI). Copy A encompasses the *B. subtilis yaaO* (Sekowska *et al.,* 1998), while copy B contains the *B. subtilis speA* gene coding for an arginine decarboxylase (NP_389346.1) (Burrell *et al.,* 2010). Two other *Firmicutes* proteins corresponding to copy B sequences (from *Clostridium difficile* YP_001087362.1 and *S. ruminantium* WP_014425426.1) (Fig. 1A) were also shown to possess an arginine decarboxylase activity (Liao *et al.,* 2008, Burrell *et al.,* 2010), which led to a hypothesis that copy B AAT-fold decarboxylases would be Adcs (Burrell *et al.,* 2010). The function of copy A awaits further investigation and, to our knowledge, the only study focused on the *yaaO* gene (copy A) concluded that in *B. subtilis* this protein had no effect on polyamine production (Sekowska *et al.,* 1998).

Regarding the evolutionary history of the wing domain-containing LAOdcs, our analysis reveals that proteobacterial LAOdcs form two monophyletic groups, Ode and LAdc. Thus, lysine and arginine decarboxylases appear more closely related to each other than to ornithine decarboxylases, although the relationships among the two Ldc families (LdcI/C and LdcA) and the Adc family are not resolved (PP <0.5) (Fig. 3A). Furthermore, we showed that the proteobacterial biosynthetic OdcC and LdcC emerged from inducible Ode and Ldc *(i.e.* Odcl and Lcdl, respectively) through two independent gene duplication events that occurred in *Enterobacteriaceae,* after the divergence of *Sodalis.* Given that both duplication events seem to occur in the same branch of the phylogenetic tree, it is tempting to hypothesize that they are linked, and that a functional connection between both biosynthetic subgroups may have existed. The emergence of biosynthetic enzymes from inducible ones may appear contra-intuitive at the first glance, but may reflect an expansion and a diversification of polyamine functions in these lineages. This would also explain why *Enterobacteriaceae* possess also a constitutive pathway of polyamine synthesis through an AR-fold Adc.

The exhaustive phylogenetic analysis of AAT-fold decarboxylases discloses multiple cases of horizontal gene transfers (e.g. within Cluster III, from *Firmicutes* to *Firmicutes* and to *Actinobacteria* within Cluster I, but also from *Proteobacteria* to *Firmicutes* within cluster II). Interestingly, the two wing-less AAT-fold LAOdc coding genes found in *L. saerimneri* 30a were proposed to result from the horizontal gene transfer of an acid stress inducible Ode from *Enterobacteriaceae,* followed by a gene duplication event (Romano *et al.,* 2013, Romano *et al.,* 2014). One of the two resulting paralogues is thought to have kept the original function (<u>WP 009553942.1</u>), while the other acquired the capacity to use lysine as substrate (<u>EKW98991.1</u>). Instead, our analysis points to a different scenario. In fact, the two copies present in *L. saerimneri* branch in two different parts of the Odd tree (Fig. S5), meaning that they have different origins, and result likely from two independent horizontal gene transfer events of Odd coding genes, whereupon one of these two acquired genes shifted secondarily towards the capacity to use lysine instead of ornithine. Interestingly, the lysine decarboxylating copy is present in various *Lactobacilli,* while the copy using ornithine as substrate is specific to *L. saerimneri* (Fig. S5), suggesting that the former was acquired first. Remarkably, these two enzymes rely on the same antiporter capable to exchange both ornithine/Put and lysine/Cad pairs, resulting in a unique three-component decarboxylation system involved in acid stress response (Romano *et al.,* 2013). This case of substrate specificity shift is probably not an exception, as exemplified by *Burkholderia* sp. AIU 395 in which an AAT-fold enzyme using lysine as substrate branches within the Adc cluster in phylogenetic trees (Fig. 3). Beside substrate shift, existence of dual specificity has been most extensively documented in the case of AR-fold LOdcs, exemplified by bifunctional enzymes of *S. ruminantium* (Takatsuka *et al.,* 2000) and *V. vulnificus* (Lee *et al.,* 2007). In particular, crystal structures of the *V. vulnificus* enzyme in complex with either Put or Cad revealed that the dual substrate specificity is based on a bridging water molecule necessary for the binding of a shorter Put ligand in addition to the longer Cad. A similar dual substrate specificity mechanism may also exist in the case of the AAT-fold LAOdc enzymes although it has not been documented up to now.

Altogether, our data indicate that functional changes affecting gene regulation, substrate fixation, and cellular function occurred several times during the evolution of AAT-fold enzymes. In the light of these observations, one may wonder if the so-called Adc, LdcC, Ldcl, OdcC, and Odd clusters defined according to phylogenetic criteria correspond indeed to homogeneous functional groups and thus if phylogenetic criteria/sequence similarity based measures are good predictors of the function of the AAT-fold enzymes. The very restricted number of experimentally characterized enzymes calls for caution and for the urgent need for additional experimental data.

One of the main results of the presented phylogenetic analysis is the identification of a novel large family of decarboxylases, called LdcA, ancestral in *Betaproteobacteria* and in *Pseudomonadaceae.* Thus, the second part of this work was dedicated to functional characterization of the LdcA from a major human opportunistic pathogen *P. aeruginosa.*

The *IdcA* gene belongs to the core genome of *P. aeruginosa,* comprising at least 4,000 conserved genes (Hilker *et al.,* 2015, Valot *et al.,* 2015). Similarly to *ldcI,* in *P. aeruginosa, Pseudomonas resinovorans, Pseudomonas denitrificans* and *Pseudomonas knackmusii, IdcA* is organized in a gene cluster with a gene coding for a homologue of the CadB antiporter, although *cadB (PA1819* in *P. aeruginosa)* appears downstream, and not upstream, of the lysine decarboxylase-encoding gene. The presence of a dedicated Lys/Cad transporter could be important from a physiological standpoint because CadB is involved not only in substrate/product exchange but also in the generation of proton motive force (Soksawatmaekhin *et al.,* 2004). Remarkably, the proximity of *cadB* and *ldcA* is an exception in the *IdcA* subfamily (Fig. S2), which does not rule out a hypothesis that LdcA could have another function in other taxa. The whole of our data suggests that Ldcl and *P. aeruginosa* LdcA are different in terms of regulation and function. Indeed, we clearly show that LdcA is not involved in acid or oxidative stress response, and its expression is triggered by neither of these stresses; instead it is controlled by ArgR and QS. More similar to LdcC, *P. aeruginosa* LdcA, and certainly the other members of this novel lysine decarboxylase subfamily, are biosynthetic enzymes responsible for the Cad production by the bacterium.

Importantly, LdcA is the only Cad producing enzyme in *P. aeruginosa* PAO1, as demonstrated by the measurement of the intracellular contents of Cad (Fig. 6). This conclusion was unexpected considering that the product of another gene, *PA4115,* was previously reported to be responsible for 25% of Cad production in overnight cultures grown in the same medium as the one used in our assays, and therefore proposed to be an Ldc (Manuel *et al.,* 2010). After carboxypenicillin treatment, the *PA4115* mutant exhibited an increased number of persisters that was significantly reduced upon addition of exogenous Cad, further supporting the hypothesis of PA4115 being an Ldc (Manuel *et al.,* 2010). Yet, our careful inspection of its sequence revealed that PA4115 belongs to the family of Lonely Guy (LOG) proteins because it contains a highly conserved PGGxGTxxE motif and a nucleotide-binding Rossmann fold. Remarkably, LOG proteins were shown to be often mis-annotated as lysine decarboxylase enzymes, yet without support by biochemical or functional data, whereas they actually possess a cytokinin-specific phosphoribohydrolase activity (Dzurova *et al.,* 2015, Seo & Kim, 2017) or a pyrimidine/purine nucleotide 5’-monophosphate nucleosidase activity (Sevin *et al.,* 2017). Recently, PPnN (or YghD) from *E. coli,* a close homologue sharing 56% identity with PA4115, has been shown to catalyze the hydrolysis of N-glycosidic bond of AMP, GMP, IMP, CMP, dTMP and UMP to form ribose 5-phosphate and the corresponding free base. Hence, it is quite likely that PA4115 catalyzes the same reaction and plays a role in maintaining the nucleotide pool homeostasis by degrading excess nucleotides in *P. aeruginosa* (Sevin *et al.,* 2017). Therefore, the reduced production of Cad in *PA4115* mutant needs to be confirmed because the relationship between PA4115 and LdcA activity is not clear and could involve indirect causes.

The persistence phenotype observed in *ldcA* mutants is consistent with the beneficial effect of LdcA on growth fitness. Indeed, recent research on bacterial persistence uncovered intracellular ATP concentrations as being one of the major factors affecting the amount of persister cells, and demonstrated that ATP levels are sufficient to predict bacterial tolerance to antibiotics (Conlon *et al.,* 2016, Shan *et al.,* 2017). Considering that Cad produced by LdcA is metabolized and used up by the Krebs cycle to create ATP and that the activity of the lysine/Cad antiporter generates proton motive force essential for ATP synthesis (Soksawatmaekhin *et al.,* 2004), one would expect that in *P. aeruginosa* the *ldcA* mutation may lead to a decrease in ATP levels, which in turn would result in an increase of persisters’ population. This hypothesis could be challenged by blocking the Cad degradation pathway or the CadB antiporter activity.

The capacity of Put and Spd to complement the growth phenotype of the *IdcA* mutant and boost *P. aeruginosa* cultures in minimal medium suggests that in *P. aeruginosa* the three major polyamines share certain properties. In the same lines, Cad was shown to substitute for Put and Spd as growth factors in *E. coli* cells depleted of these two polyamines (Igarashi *et al.,* 1986). The growth phenotype described in our work is observed under specific growth conditions. It reveals the importance of Cad when growing *P. aeruginosa* in minimal medium and highlights a certain redundancy in the function of polyamines. It remains to be investigated whether the phenotype is linked to a regulatory effect of Cad or to its anabolism. Recent literature about *Eikenella corrodens* has pointed out the importance of an AAT-fold Ldc (belonging to Ldcl group) as a virulence factor against eukaryotic cells that acts through depletion of essential lysine (Lohinai *et al.,* 2015). Therefore the potential role of LdcA in the virulence of *P. aeruginosa* may warrant further investigation. In the present study, we observed that the absence of the enzyme did not affect T3SS-dependent cytotoxicity or the mobility relying on flagellum and Type IV pili (not shown). However, it would also be relevant to probe the importance of LdcA during mouse infection, where the proper functioning of *P. aeruginosa* metabolism is primordial for virulence.

Our study reveals that the *ldcA* gene is relatively ancient in *Proteobacteria,* being ancestral in *Betaproteobacteria* and in *Pseudomonadaceae,* two taxa that cover a wide range of ecological niches. Information about regulation and function of an enzyme of the previously unknown LdcA subfamily enables a step further towards understanding of the evolution and the importance of Cad metabolism in bacteria.

## Experimental procedures

### Phytogeny: Dataset assembly

Functionally characterized sequences of AAT-fold LAOdcs were retrieved from NCBI: Ldcl (NP_418555.1), LdcC (NP_414728.1), Adc (NP_418541.1), OdcC (NP_417440.1), and Odd (NP_415220.1) from *E coli* str. K-12 substr., MG1655 and LdcA (NP_250509.1) from *P. aeruginosa* PAOl. These sequences were used as seeds to query a local database containing 4,466 complete proteomes of prokaryotes downloaded from the NCBI (ftp://ftp.ncbi.nlm.nih.gov) with the BLASTP 2.2.6 software (Altschul *et al.,* 1997) using default parameters. Homologues of LAODc were retrieved and aligned using MAFFT v7 (Katoh & Standley, 2013). The resulting multiple alignment was used to build a HMM profile with the HMMbuild program from the HMMER v3.1b1 package (McClure MA, 1996). This profile was then used to query the local database of complete proteomes with the HMMsearch program. Sequences with e-values lower than 2.2e-13 were retrieved. Finally, the search for potential unannotated sequences was performed using TBLASTN 2.2.6 on genomic sequences corresponding to the 4,466 complete proteomes. This led to the identification of 4,090 homologous sequences.

### Phylogeny: Phylogenetic inference

To limit taxonomic redundancy for phylogenetic analyses, a sampling of the retrieved sequences by selecting randomly one strain per species has been performed. Multiple alignments were built with MAFFT using the L-INS-i option that allows the construction of accurate multiple alignments and trimmed with BMGE v1.1 with matrix substitution BLOSUM30 (Criscuolo & Gribaldo, 2010).

Maximum likelihood trees were inferred with PhyML 3.1 (Guindon *et al.,* 2010). The best suited evolutionary models were selected using the model test tool implemented in IQ-TREE v1.4.1 according to the Bayesian information criterion (BIC) (Nguyen *et al.,* 2015). The robustness of the inferred trees was assessed using the non-parametric bootstrap procedure implemented in PhyML (100 replicates of the original datasets). Bayesian trees were inferred using MrBayes v3.2.6 (Ronquist & Huelsenbeck, 2003). Two runs were launched with four chains for each run (50,000 iterations). The first 25% of the trees were discarded as burn-in and chain convergence has been checked by analysing the evolution of the Ln(L) curve and checking the average standard deviation of split frequencies values. Figures of trees have been generated using EvolView (He *et al.,* 2016) (http://nar.oxfordiournals.org/content/44/Wl/W236) and iTOL (Letunic & Bork, 2016). Genomic context figures have been generated using GeneSpy (https://lbbe.univ-lvonl.fr/GeneSpy/) developed by P.S. Garcia.

Reference phylogenies of *Firmicutes, Cyanobacteria, Proteobacteria,* and Enterobacteriaceae strains contained in our local database were inferred using ribosomal proteins as suggested elsewhere (Ramulu *et al.,* 2014). The reference tree of *Firmicutes* has been rooted according to a recent study (Antunes *et al.,* 2016). The reference tree of *Cyanobacteria* has been rooted by including ribosomal protein sequences from *Natranaerobius thermophilus (Firmicutes)* and *Streptomyces albulus (Actinobacteria).* The reference tree of *Proteobacteria* has been rooted according to Gupta (Gupta, 2000). Finally, the reference phylogeny of *Enterobacteriaceae* has been rooted using with *Shewanella baltica (Alteromonadales)* and *Pasteurella multocida (Pasteurellales)* ribosomal protein sequences.

For each analysis, the ribosomal protein sequences were identified using the RiboDB database engine (Jauffrit *et al.,* 2016) and aligned with MAFFT using the L-INS-i option. The resulting multiple alignments were trimmed as described above and combined to build a large supermatrix that was used to build maximum likelihood phylogenetic trees.

### Bacterial strains, plasmids, and growth conditions

The bacterial strains and plasmids used in this study are listed in Table SI in the supplemental material. Bacteria were cultivated aerobically at 37°C in rich Lysogeny Broth (LB) medium, in Mueller Hinton II (Becton Dickinson) or in minimal medium P (30mM Na_2_HPO_4_, 14 mM KH_2_PO_4_, 20mM (NH4)_2_SO_4_, 1 mM MgSO_4_, 4μM FeSO_4_, 0.4 μM Pyridoxal-5’-phosphate, pH 7.4) (Haas *et al.,* 1977) containing the indicated carbon and nitrogen sources. *P. aeruginosa* was also cultured on Pseudomonas Isolation Agar plates (PIA; Difco). When required, antibiotics were added at the following concentrations (in μg/ml): 100 (ampicillin), 25 (gentamycin), 25 (kanamycin) and 10 (tetracycline) for *E. coli,* 200 (carbenicillin), 200 (gentamycin) and 200 (tetracycline) for *P. aeruginosa.*

### Genetic manipulations

To delete *IdcA* gene, fused uspstream and downstream flanking regions of the gene were generated by “Splicing by Overlap Extension” (SOE)-PCR using PAO1 genomic DNA as matrix and appropriate primer pairs. The resulting fragment of 819 bp was cloned into pCR-Blunt ll-TOPO vector, sequenced and then subcloned into *BamHI-HindIII* sites of pEXG2, leading to pEXG2Δ*ldcA.* The suicide plasmid carries the counter-selectable *sacB* marker from *B. subtilis* which confers sensitivity to sucrose. The plasmid was mobilized into *P. aeruginosa* strain by triparental mating, using the conjugative properties of the helper plasmid pRK2013. Co-integration events were selected on PIA plates containing gentamycin. Single colonies were then plated on PIA medium containing 5% (w/v) sucrose to select for the loss of plasmid: the resulting sucrose-resistant strains were checked for antibiotic sensitivity and for *ldcA* (wild-type or truncated gene) genotype by PCR.

To complement the *ldcA* mutant, a 2785 bp-long fragment encompassing the coding sequence and 495 bases upstream the ATG was PCR amplified from PAO1 genomic DNA using appropriate primer pairs. The PCR product was cloned into pCR-Blunt ll-TOPO and sequenced. The *Spe*I restriction fragment was subcloned in mini-CTXl cut with the same enzyme, leading to *miniCTX-IdcA.* A mini-CTX derivative was used to construct a *lacZ* reporter vector. PCR amplification was used to produce the 548 bp-long *ldcA* promoter fragment (−498/+44 relative to translation initiation) with appropriate primers. After ligation into pCR-Blunt ll-TOPO vector and sequencing, the fragment was sub-cloned into the *Xhol-EcoRl* sites of mini-CTX-lacZ. Both *miniCTX-ldcA* and mini-CTX-*PldcA-lacZ* were introduced into *P. aeruginosa* by triparental conjugation and the transconjugants were selected on PIA plates containing tetracycline. The pFLP2 plasmid was then used to excise the FRT cassette as described (Hoang *et al.,* 1998).

Plasmids and primers used in PCR are listed in the Tables SI and S2, respectively.

### β-Galactosidase assays

Bacteria were grown aerobically at 37°C in 100 ml flasks with agitation (300 rpm). At the indicated OD, (β-galactosidase activity was assayed as described (Miller, 1972), with details reported in (Thibault *et al.,* 2009).

### Intracellular metabolite analysis

Strains were first isolated from an overnight solid culture at 37°C on Mueller Hinton II agar 1.5%. Precultures and cultures were performed on a Minitron II rotary shaker at 220 rpm (Infors HT) under aerobic conditions (10% of total volume of Erlenmeyer flask). A few bacterial colonies from the agar plate were precultured overnight at 37°C and an aliquot was withdrawn and diluted to an OD_600_ of ∼0.1 in a fresh culture medium for the culture step. Growth curves were determined for each strain and used to determine their respective concentration (CFU/ml) and the OD at which the bacteria should be harvested to correspond to early-, mid-, and late-exponential phases.

The protocol of the intracellular metabolites sampling was adapted from a previously described procedure (Aros-Calt *et al.,* 2015). In brief, a 5 ml aliquot of cell culture broth was taken from the main culture and was filtered in a few seconds using poly(ether sulfone) sterile membrane disc filters (Supor450, 0.45 μm pore size, PALL) mounted on a Millipore filtration device (Darmstadt). The bacteria on the filter were quickly washed with 5 ml of 0.6% NaCI solution maintained at room temperature. The filter was then rapidly transferred to a 50 ml Falcon tube containing 5 ml of cold 60% ethanol (v/v, ≤ −20°C). The Falcon tube was subsequently quickly immersed in liquid nitrogen. Following this quenching step, tubes containing bacteria on filters in the extraction solution were vortexed 10 times on ice to remove the cells from the filter. Then, a 1 ml aliquot of the cell suspension was transferred to 2 ml tubes containing 0.1 mm glass beads (Bertin Technologies). Cell disruption was performed by three cycles in a Precellys 24 homogenizer (Bertin Technologies) for 30 s at 3,800 rpm at ∼4°C. The glass beads and cell debris were separated from the supernatant by centrifugation for 5 min at 4°C and 10,000g. A 400 μL aliquot of the supernatant was withdrawn and further vacuum-dried using a SpeedVac instrument (Thermo Fisher Scientific) and stored at −80°C until analysis. Dried extracts were dissolved in an adjusted volume of 95% mobile phase A / 5% mobile phase B to obtain the equivalent of 1.25 10^7^ CFU in 15μl before analysis by LC/HR-MS.

To detect intracellular metabolites, LC/HR-MS experiments were performed using a Dionex Ultimate chromatographic system (Thermo Fisher Scientific) coupled to an Exactive (Orbitrap) mass spectrometer from Thermo Fisher Scientific fitted with an electrospray source. The mass spectrometer was externally calibrated before each analysis using the manufacturer’s predefined methods and recommended calibration mixture provided by the manufacturer. Chromatographic separation was performed on a Discovery HS F5 PFPP 5 μm, 2.1 ×; 250 mm column (Sigma) at 30°C. The chromatographic system was equipped with an on-line pre-filter (Thermo Fisher Scientifics). Mobile phases were 100% water (A) and 100% ACN (B), both of which containing 0.1% formic acid. Chromatographic elution was achieved with a flow rate of 250 μl/mm. After injection 15 μl of sample, elution started with an isocratic step of 2 min at 5% phase B, followed by a linear gradient from 5 to 100% of phase B in 18 min. These proportions were kept constant for 4 min before returning to 5% of phase B and letting the system equilibrate for 6 min. The column effluent was directly introduced into the electrospray source of the mass spectrometer, and analyses were performed in the positive ion mode. Source parameters were as follows: capillary voltage set at 5 kV; capillary temperature at 300°C; sheath and auxiliary gas (nitrogen) flow rates at 50 and 25 arbitrary units, respectively; mass resolution power of the analyzer set at 50,000 at m/z 200 (full width at half maximum, FWHM) for singly charged ions. The acquisition was achieved from m/z 50 to 250 in the positive ionization mode during the first 12 min of the run. Under these conditions were achieved a good separation and detection (with an average mass accuracy better than 3ppm) of the targeted molecules (under their [M+H]^+^ form). These species were readily identified in the extracts through the use of the corresponding commercial molecules obtained from Sigma-Aldrich. Extracted ion chromatograms were generated and resulting peaks integrated using the Xcalibur software (version 2.1, Thermo Fisher Scientific) for Spd ([M+H]^+^ at theoretical m/z 146.1652, retention time 5.24 min), Put (m/z 89.1073, 3.63 min) and Cad (m/z 103.1230, 3.94 min).

## Acknowledgments

We thank Rémi Peyraud and Ina Attrée for helpful discussions. This work has received funding from GRAL (ANR-10-LABX-49-01), the European Union’s Horizon 2020 research and innovation programme under grant agreement No 647784, and the ANR-16-CE02-0005. This work was also supported by CEA and the French Ministry of Research and National Research Agency as part of the French metabolomics and fluxomics infrastructure (MetaboHUB, ANR-ll-INBS-0010 grant). Diego Carriel was supported by a PhD grant from LABEX GRAL and Pierre S. Garcia was supported by a grant from the ARC Santé from the Region Auvergne Rhône-Alpes.

## Author contributions

DCL, PSG, FC, and PL performed experiments. All authors analyzed data. IG, SE and CBA designed and supervised the overall study and wrote the paper with contribution from all authors.

## Abbreviated Summary

Bacterial polyamines are involved in many fundamental processes and are mainly synthetized by dedicated lysine, arginine and ornithine decarboxylases or LAOdcs. Our exhaustive phylogenetic analysis reveals evolutionary history of LAOdcs and discloses a hitherto overlooked LdcA subfamily. We show that LdcA is playing an important role in growth and persistence of the major multidrug resistant human pathogen *P. aeruginosa,* exerted through cadaverine biosynthesis and concomitant regulation of intracellular levels of putrescine and spermidine.

## Supporting informations

### Supplemental material and methods

**Fig. SI.** AAT-fold LAOdc are widespread in *Bacteria*

(A) Taxonomic distribution of AAT-fold LAOdc in prokaryotes. For each phylum and class, ratio corresponds to the number of proteomes containing at least one homologue on the number of proteomes present in our database. Taxa represented by at least 10 proteomes, among which more than 20% contained at least one AAT-fold LAOdc homologue are highlighted by colors.

(B) Unrooted ML tree of AAT-fold LAOdcs (PhyML, LG+I+G4, 1,117 sequences, 273 amino-acid positions). The tree can be divided in three parts: Cluster I corresponds to wing-less LAOdc mainly from *Firmicutes, Actinobacteria* and *Cyanobacteria;* Cluster II encompasses wing domain containing LAOdc mainly from *Proteobacteria,* while Cluster III gathers wing-less and wing domain-containing LAOdc from various and unrelated taxa. The scale bar represents the number of substitutions per site. Bootstrap values (BV) associated to branches separating the three clusters are indicated.

**Fig. S2.** Genomic context of LAOdc genes from Cluster II

Genomic context of LdcC, Ldcl, OdcC, Odd, Adc, and Ladc coding genes in a sample of Proteobacteria. Black arrows: LAOdc coding genes, colored arrows: conserved neighbor genes.

**FIG S3** Influence of stress on *ldcA* expression and growth fitness of *P. aeruginosa.* (A) Effect of acid and oxidative stress during growth in rich medium. At T_0_, HCI was added to shift the pH of LB medium from neutral to pH 5 while H_2_O_2_ was added to a final entration of 1mM. β-galactosidase activity of PAO1:*:PldcA-lacZ* strain was measured at T_0_, then monitored 30 and 60 min after treatment. B) Growth of PAO1 in minimal medium at different pHs was evaluated using Biolog high-throughput system. Each point corresponds to the “Area Under the Curve” (AUC) measured after 24h of bacterial growth.

**FIG S4** LdcA function affects carbenicillin persistence. Percentage of survivors in rich medium (cation-adjusted Mueller Hinton Broth) after 24h of carbenicillin treatment. Growth was performed in erlenmeyer flasks at 37°C with agitation (300 RPM). Percentage of survivors was calculated from CFU counting after 24h of antibiotic treatment at 500μg/ml (8X MIC).

**FIG. S5** Maximum likelihood tree of Odcl/OdcC sequences (in blue) (PhyML, LG+I+G4, 514 sequences, 583 amino-acid positions), disclosing the proteobacterial origin of the Odd sequences reported by previous studies in a few firmicutes (in dark red) (Romano *et al.,* 2013, Romano *et al.,* 2014). These sequences were likely acquired through at least three horizontal gene transfers indicated by dark yellow branches. The tree was rooted with Adc (orange triangle), LdcA (green triangle), and Ldcl/C (blue triangle) sequences. The pink triangle correspond to OdcC sequences. Numbers at branches correspond to aLRT supports. The scale bar indicates the average number of substitutions per site.

**Table SI** List of bacterial strains and plasmids used in this work.

**Table S2** Primers used in this study.

## REFERENCES

Altschul, S.F., T.L. Madden, A.A. Schaffer, J. Zhang, Z. Zhang, W. Miller & D.J. Lipman, (1997) Gapped BLAST and PSI-BLAST: a new generation of protein database search programs. Nucleic Acids Res 25: 3389–3402.

Antunes, L.C., D. Poppleton, A. Klingl, A. Criscuolo, B. Dupuy, C. Brochier-Armanet, C. Beloin & S. Gribaldo, (2016) Phylogenomic analysis supports the ancestral presence of LPS-outer membranes in the Firmicutes. Elife 5.

Aros-Calt, S., B.H. Muller, S. Boudah, C. Ducruix, G. Gervasi, C. Junot & F. Fenaille, (2015) Annotation of the Staphylococcus aureus Metabolome Using Liquid Chromatography Coupled to High-Resolution Mass Spectrometry and Application to the Study of Methicillin Resistance. J Proteome Res 14: 4863–4875.

Bodey, G.P., R. Bolivar, V. Fainstein & L. Jadeja, (1983) Infections caused by Pseudomonas aeruginosa. Rev Infect Dis 5: 279–313.

Bunsupa, S., K. Katayama, E. Ikeura, A. Oikawa, K. Toyooka, K. Saito & M. Yamazaki, (2012) Lysine decarboxylase catalyzes the first step of quinolizidine alkaloid biosynthesis and coevolved with alkaloid production in leguminosae. Plant Cell 24: 1202–1216.

Burrell, M., C.C. Hanfrey, L.N. Kinch, K.A. Elliott & A.J. Michael, (2012) Evolution of a novel lysine decarboxylase in siderophore biosynthesis. Mol Microbiol 86: 485–499.

Burrell, M., C.C. Hanfrey, E.J. Murray, N.R. Stanley-Wall & A.J. Michael, (2010) Evolution and multiplicity of arginine decarboxylases in polyamine biosynthesis and essential role in Bacillus subtilis biofilm formation. J Biol Chem 285: 39224–39238.

Chou, H.T., M. Hegazy & C.D. Lu, (2010) L-lysine catabolism is controlled by L-arginine and ArgR in Pseudomonas aeruginosa PAO1. J Bacteriol 192: 5874–5880.

Conlon, B.P., S.E. Rowe, A.B. Gandt, A.S. Nuxoll, N.P. Donegan, E.A. Zalis, G. Clair, J.N. Adkins, A.L. Cheung & K. Lewis, (2016) Persister formation in Staphylococcus aureus is associated with ATP depletion. Nat Microbiol 1: 16051.

Criscuolo, A. & S. Gribaldo, (2010) BMGE (Block Mapping and Gathering with Entropy): a new software for selection of phylogenetic informative regions from multiple sequence alignments. BMC Evol Biol 10: 210.

Dela Vega, A.L. & A.H. Delcour, (1996) Polyamines decrease Escherichia coli outer membrane permeability. J Bacteriol 178: 3715–3721.

Di Martino, M.L., R. Campilongo, M. Casalino, G. Micheli, B. Colonna & G. Prosseda, (2013) Polyamines: emerging players in bacteria-host interactions. Int J Med Microbiol 303: 484–491.

Dzurova, L., F. Forneris, S. Savino, P. Galuszka, J. Vrabka & I. Frebort, (2015) The three-dimensional structure of “Lonely Guy” from Claviceps purpurea provides insights into the phosphoribohydrolase function of Rossmann fold-containing lysine decarboxylase-like proteins. Proteins 83: 1539–1546.

El Bakkouri, M., I. Gutsche, U. Kanjee, B. Zhao, M. Yu, G. Goret, G. Schoehn, W.P. Burmeister & W.A. Houry, (2010) Structure of RavA MoxR AAA+ protein reveals the design principles of a molecular cage modulating the inducible lysine decarboxylase activity. Proc Natl Acad Sci U S A 107: 22499–22504.

Eliot, A.C. & J.F. Kirsch, (2004) Pyridoxal phosphate enzymes: mechanistic, structural, and evolutionary considerations. Annu Rev Biochem 73: 383–415.

Fothergill, J.C. & J.R. Guest, (1977) Catabolism of L-lysine by Pseudomonas aeruginosa. J Gen Microbiol 99: 139–155.

Fritz, G., C. Koller, K. Burdack, L. Tetsch, I. Haneburger, K. Jung & U. Gerland, (2009) Induction kinetics of a conditional pH stress response system in Escherichia coli. J Mol Biol 393: 272–286.

Gale, E.F. & H.M. Epps, (1944) Studies on bacterial amino-acid decarboxylases: 1. l(+)-lysine decarboxylase. Biochem J 38: 232–242.

Gale, E.F. & W.E. Van Heyningen, (1942) The effect of the pH and the presence of glucose during growth on the production of alpha and theta toxins and hyaluronidase by Clostridium welchii. Biochem J 36: 624–630.

Gellatly, S.L. & R.E. Hancock, (2013) Pseudomonas aeruginosa: new insights into pathogenesis and host defenses. Pathog Dis 67: 159–173.

Guindon, S., J.F. Dufayard, V. Lefort, M. Anisimova, W. Hordijk & O. Gascuel, (2010) New algorithms and methods to estimate maximum-likelihood phylogenies: assessing the performance of PhyML 3.0. Syst Biol 59: 307–321.

Gupta, R.S., (2000) The phylogeny of proteobacteria: relationships to other eubacterial phyla and eukaryotes. FEMS Microbiol Rev 24: 367–402.

Haas, D., B.W. Holloway, A. Schambock & T. Leisinger, (1977) The genetic organization of arginine biosynthesis in Pseudomonas aeruginosa. Mol Gen Genet 154: 7–22.

He, Z., H. Zhang, S. Gao, M.J. Lercher, W.H. Chen & S. Hu, (2016) Evolview v2: an online visualization and management tool for customized and annotated phylogenetic trees. Nucleic Acids Res 44: W236–241.

Hilker, R., A. Munder, J. Klockgether, P.M. Losada, P. Chouvarine, N. Cramer, C.F. Davenport, S. Dethlefsen, S. Fischer, H. Peng, T. Schonfelder, O. Turk, L. Wiehlmann, F. Wolbeling, E. Gulbins, A. Goesmann & B. Tummler, (2015) Interclonal gradient of virulence in the Pseudomonas aeruginosa pangenome from disease and environment. Environ Microbiol 17: 29–46.

Hoang, T.T., R.R. Karkhoff-Schweizer, A.J. Kutchma & H.P. Schweizer, (1998) A broad-host-range Flp-FRT recombination system for site-specific excision of chromosomally-located DNA sequences: application for isolation of unmarked Pseudomonas aeruginosa mutants. Gene 212: 77–86.

Igarashi, K. & K. Kashiwagi, (2006) Polyamine Modulon in Escherichia coli: genes involved in the stimulation of cell growth by polyamines. J Biochem 139: 11–16.

Igarashi, K. & K. Kashiwagi, (2015) Modulation of protein synthesis by polyamines. IUBMB Life 67: 160–169.

Igarashi, K., K. Kashiwagi, H. Hamasaki, A. Miura, T. Kakegawa, S. Hirose & S. Matsuzaki, (1986) Formation of a compensatory polyamine by Escherichia coli polyamine-requiring mutants during growth in the absence of polyamines. J Bacteriol 166: 128–134.

Jauffrit, F., S. Penel, S. Delmotte, C. Rey, D.M. de Vienne, M. Gouy, J.P. Charrier, J.P. Flandrois & C. Brochier-Armanet, (2016) RiboDB Database: A Comprehensive Resource for Prokaryotic Systematics. Mol Biol Evol 33: 2170–2172.

Kamio, Y. & K. Nakamura, (1987) Putrescine and cadaverine are constituents of peptidoglycan in Veillonella alcalescens and Veillonella parvula. J Bacteriol 169: 2881–2884.

Kamio, Y., H. Poso, Y. Terawaki & L. Paulin, (1986) Cadaverine covalently linked to a peptidoglycan is an essential constituent of the peptidoglycan necessary for the normal growth in Selenomonas ruminantium. J Biol Chem 261: 6585–6589.

Kandiah, E., D. Carriel, J. Perard, H. Malet, M. Bacia, K. Liu, S.W. Chan, W.A. Houry, S. Ollagnier de Choudens, S. Elsen & I. Gutsche, (2016) Structural insights into the Escherichia coli lysine decarboxylases and molecular determinants of interaction with the AAA+ ATPase RavA. Sci Rep 6: 24601.

Kang, I.H., J.S. Kim, E.J. Kim & J.K. Lee, (2007) Cadaverine protects Vibrio vulnificus from superoxide stress. J Microbiol Biotechnol 17: 176–179.

Kanjee, U., I. Gutsche, E. Alexopoulos, B. Zhao, M. El Bakkouri, G. Thibault, K. Liu, S. Ramachandran, J. Snider, E.F. Pai & W.A. Houry, (2011a) Linkage between the bacterial acid stress and stringent responses: the structure of the inducible lysine decarboxylase. EMBO J 30: 931–944.

Kanjee, U., I. Gutsche, S. Ramachandran & W.A. Houry, (2011b) The enzymatic activities of the Escherichia coli basic aliphatic amino acid decarboxylases exhibit a pH zone of inhibition. Biochemistry 50: 9388–9398.

Kanjee, U. & W.A. Houry, (2013) Mechanisms of acid resistance in Escherichia coli. Annu Rev Microbiol 67: 65–81.

Karatan, E. & A.J. Michael, (2013) A wider role for polyamines in biofilm formation. Biotechnol Lett 35: 1715–1717.

Katoh, K. & D.M. Standley, (2013) MAFFT multiple sequence alignment software version 7: improvements in performance and usability. Mol Biol Evol 30: 772–780.

Kikuchi, Y., O. Kurahashi, T. Nagano & Y. Kamio, (1998) RpoS-dependent expression of the second lysine decarboxylase gene in Escherichia coli. Biosci Biotechnol Biochem 62: 1267–1270.

Kim, J.S., S.H. Choi & J.K. Lee, (2006) Lysine decarboxylase expression by Vibrio vulnificus is induced by SoxR in response to superoxide stress. J Bacteriol 188: 8586–8592.

Kim, S.H., Y. Wang, M. Khomutov, A. Khomutov, C. Fuqua & A.J. Michael, (2016) The Essential Role of Spermidine in Growth of Agrobacterium tumefaciens Is Determined by the 1,3-Diaminopropane Moiety. ACS Chem Biol 11: 491–499.

Kuper, C. & K. Jung, (2005) CadC-mediated activation of the cadBA promoter in Escherichia coli. J Mol Microbiol Biotechnol 10: 26–39.

Kwon, D.H. & C.D. Lu, (2006) Polyamines induce resistance to cationic peptide, aminoglycoside, and quinolone antibiotics in Pseudomonas aeruginosa PAO1. Antimicrob Agents Chemother 50: 1615–1622.

Lee, J., A.J. Michael, D. Martynowski, E.J. Goldsmith & M.A. Phillips, (2007) Phylogenetic diversity and the structural basis of substrate specificity in the beta/alpha-barrel fold basic amino acid decarboxylases. J Biol Chem 282: 27115–27125.

Lee, Y.S. & Y.D. Cho, (2001) Identification of essential active-site residues in ornithine decarboxylase of Nicotiana glutinosa decarboxylating both L-ornithine and L-lysine. Biochem J 360: 657–665.

Letunic, I. & P. Bork, (2016) Interactive tree of life (iTOL) v3: an online tool for the display and annotation of phylogenetic and other trees. Nucleic Acids Res 44: W242–245.

Liao, S., P. Poonpairoj, K.C. Ko, Y. Takatuska, Y. Yamaguchi, N. Abe, J. Kaneko & Y. Kamio, (2008) Occurrence of agmatine pathway for putrescine synthesis in Selenomonas ruminatium. Biosci Biotechnol Biochem 72: 445–455.

Lightfoot, H.L. & J. Hall, (2014) Endogenous polyamine function--the RNA perspective. Nucleic Acids Res 42: 11275–11290.

Lohinai, Z., B. Keremi, E. Szoko, T. Tabi, C. Szabo, Z. Tulassay, J.C. DiCesare, C.A. Davis, L.M. Collins & M. Levine, (2015) Biofilm Lysine Decarboxylase, a New Therapeutic Target for Periodontal Inflammation. J Periodontol 86: 1176–1184.

Lu, C.D., Z. Yang & W. Li, (2004) Transcriptome analysis of the ArgR regulon in Pseudomonas aeruginosa. J Bacteriol 186: 3855–3861.

Madhuri Indurthi, S., H.T. Chou & C.D. Lu, (2016) Molecular characterization of lysR-lysXE, gcdR-gcdHG and amaR-amaAB operons for lysine export and catabolism: a comprehensive lysine catabolic network in Pseudomonas aeruginosa PAO1. Microbiology 162: 876–888.

Malet, H., K. Liu, M. El Bakkouri, S.W. Chan, G. Effantin, M. Bacia, W.A. Houry & I. Gutsche, (2014) Assembly principles of a unique cage formed by hexameric and decameric E. coli proteins. Elife 3: e03653.

Manuel, J., G.G. Zhanel & T. de Kievit, (2010) Cadaverine suppresses persistence to carboxypenicillins in Pseudomonas aeruginosa PAO1. Antimicrob Agents Chemother 54: 5173–5179.

McClure MA, S.C., Elton P., (1996) Parameterization studies for the SAM and HMMER methods of hidden markov model generation. In: Proceedings / International Conference on Intelligent Systems for Molecular Biology. pp.

Merrell, D.S. & A. Camilli, (2000) Regulation of vibrio cholerae genes required for acid tolerance by a member of the “ToxR-like” family of transcriptional regulators. J Bacteriol 182: 5342–5350.

Michael, A.J., (2016a) Biosynthesis of polyamines and polyamine-containing molecules. Biochem J 473: 2315–2329.

Michael, A.J., (2016b) Polyamines in Eukaryotes, Bacteria, and Archaea. J Biol Chem 291: 14896–14903.

Miller, J., (1972) Experiments in Molecular Genetics. In. C.S.H.L. Press (ed). Cold Spring Harbor, pp.

Nguyen, L.T., H.A. Schmidt, A. von Haeseler & B.Q. Minh, (2015) IQ-TREE: a fast and effective stochastic algorithm for estimating maximum-likelihood phylogenies. Mol Biol Evol 32: 268–274.

Ochsner, U.A., M.L. Vasil, E. Alsabbagh, K. Parvatiyar & D.J. Hassett, (2000) Role of the Pseudomonas aeruginosa oxyR-recG operon in oxidative stress defense and DNA repair: OxyR-dependent regulation of katB-ankB, ahpB, and ahpC-ahpF. J Bacteriol 182: 4533–4544.

Pezzulo, A.A., X.X. Tang, M.J. Hoegger, M.H. Abou Alaiwa, S. Ramachandran, T.O. Moninger, P.H. Karp, C.L. Wohlford-Lenane, H.P. Haagsman, M. van Eijk, B. Banfi, A.R. Horswill, D.A. Stoltz, P.B. McCray, Jr., M.J. Welsh & J. Zabner, (2012) Reduced airway surface pH impairs bacterial killing in the porcine cystic fibrosis lung. Nature 487: 109–113.

Rahman, M. & P.H. Clarke, (1980) Genes and enzymes of lysine catabolism in Pseudomonas aeruginosa. J Gen Microbiol 116: 357–369.

Ramulu, H.G., M. Groussin, E. Talla, R. Planel, V. Daubin & C. Brochier-Armanet, (2014) Ribosomal proteins: toward a next generation standard for prokaryotic systematics? Mol Phylogenet Evol 75: 103–117.

Romano, A., V. Ladero, M.A. Alvarez & P.M. Lucas, (2014) Putrescine production via the ornithine decarboxylation pathway improves the acid stress survival of Lactobacillus brevis and is part of a horizontally transferred acid resistance locus. Int J Food Microbiol 175: 14–19.

Romano, A., H. Trip, J.S. Lolkema & P.M. Lucas, (2013) Three-component lysine/ornithine decarboxylation system in Lactobacillus saerimneri 30a. J Bacteriol 195: 1249–1254.

Ronquist, F. & J.P. Huelsenbeck, (2003) MrBayes 3: Bayesian phylogenetic inference under mixed models. Bioinformatics 19: 1572–1574.

Schuster, M., A.C. Hawkins, C.S. Harwood & E.P. Greenberg, (2004) The Pseudomonas aeruginosa RpoS regulon and its relationship to quorum sensing. Mol Microbiol 51: 973–985.

Sekowska, A., P. Bertin & A. Danchin, (1998) Characterization of polyamine synthesis pathway in Bacillus subtilis 168. Mol Microbiol 29: 851–858.

Seo, H. & K.J. Kim, (2017) Structural basis for a novel type of cytokinin-activating protein. Sci Rep 7: 45985.

Sevin, D.C., T. Fuhrer, N. Zamboni & U. Sauer, (2017) Nontargeted in vitro metabolomics for high-throughput identification of novel enzymes in Escherichia coli. Nat Methods 14: 187–194.

Shah, P. & E. Swiatlo, (2008) A multifaceted role for polyamines in bacterial pathogens. Mol Microbiol 68: 4–16.

Shan, Y., A. Brown Gandt, S.E. Rowe, J.P. Deisinger, B.P. Conlon & K. Lewis, (2017) ATP-Dependent Persister Formation in Escherichia coli. MBio 8.

Snider, J., I. Gutsche, M. Lin, S. Baby, B. Cox, G. Butland, J. Greenblatt, A. Emili & W.A. Houry, (2006) Formation of a distinctive complex between the inducible bacterial lysine decarboxylase and a novel AAA+ ATPase. J Biol Chem 281: 1532–1546.

Soksawatmaekhin, W., A. Kuraishi, K. Sakata, K. Kashiwagi & K. Igarashi, (2004) Excretion and uptake of cadaverine by CadB and its physiological functions in Escherichia coli. Mol Microbiol 51: 1401–1412.

Stover, C.K., X.Q. Pham, A.L. Erwin, S.D. Mizoguchi, P. Warrener, M.J. Hickey, F.S. Brinkman, W.O. Hufnagle, D.J. Kowalik, M. Lagrou, R.L. Garber, L. Goltry, E. Tolentino, S. Westbrock-Wadman, Y. Yuan, L.L. Brody, S.N. Coulter, K.R. Folger, A. Kas, K. Larbig, R. Lim, K. Smith, D. Spencer, G.K. Wong, Z. Wu, I.T. Paulsen, J. Reizer, M.H. Saier, R.E. Hancock, S. Lory & M.V. Olson, (2000) Complete genome sequence of Pseudomonas aeruginosa PAO1, an opportunistic pathogen. Nature 406: 959–964.

Sugawara, A., D. Matsui, N. Takahashi, M. Yamada, Y. Asano & K. Isobe, (2014) Characterization of a pyridoxal-5’-phosphate-dependent l-lysine decarboxylase/oxidase from Burkholderia sp. AIU 395. J Biosci Bioeng 118: 496–501.

Tabor, C.W. & H. Tabor, (1985) Polyamines in microorganisms. Microbiol Rev 49: 81–99.

Tabor, H., E.W. Hafner & C.W. Tabor, (1980) Construction of an Escherichia coli strain unable to synthesize putrescine, spermidine, or cadaverine: characterization of two genes controlling lysine decarboxylase. J Bacteriol 144: 952–956.

Tabor, H. & C.W. Tabor, (1964) Spermidine, Spermine, and Related Amines. Pharmacol Rev 16: 245–300.

Takatsuka, Y., Y. Yamaguchi, M. Ono & Y. Kamio, (2000) Gene cloning and molecular characterization of lysine decarboxylase from Selenomonas ruminantium delineate its evolutionary relationship to ornithine decarboxylases from eukaryotes. J Bacteriol 182: 6732–6741.

Thibault, J., E. Faudry, C. Ebel, I. Attree & S. Elsen, (2009) Anti-activator ExsD forms a 1:1 complex with ExsA to inhibit transcription of type III secretion operons. J Biol Chem 284: 15762–15770.

Valot, B., C. Guyeux, J.Y. Rolland, K. Mazouzi, X. Bertrand & D. Hocquet, (2015) What It Takes to Be a Pseudomonas aeruginosa? The Core Genome of the Opportunistic Pathogen Updated. PLoS One 10: e0126468.

Viala, J.P., S. Meresse, B. Pocachard, A.A. Guilhon, L. Aussel & F. Barras, (2011) Sensing and adaptation to low pH mediated by inducible amino acid decarboxylases in Salmonella. PLoS One 6: e22397.

Zhao, B. & W.A. Houry, (2010) Acid stress response in enteropathogenic gammaproteobacteria: an aptitude for survival. Biochem Cell Biol 88: 301–314.

